# High-resolution structures map the metal import pathway in an Nramp transporter

**DOI:** 10.1101/2022.09.08.507188

**Authors:** Shamayeeta Ray, Samuel P. Berry, Eric A. Wilson, Casey H. Zhang, Mrinal Shekhar, Abhishek Singharoy, Rachelle Gaudet

**Affiliations:** Department of Molecular and Cellular Biology, Harvard University, Cambridge, MA USA; School of Molecular Sciences, Arizona State University, Tempe, AZ USA; Grossman School of Medicine, New York University, New York, NY USA; Broad Institute of MIT and Harvard, Cambridge, MA USA

## Abstract

Transporters of the Nramp (Natural resistance-associated macrophage protein) family import divalent transition metal ions into cells of most organisms. By supporting metal homeostasis, Nramps prevent disorders related to metal insufficiency or overload. Previous studies revealed that Nramps take on a LeuT fold and identified the metal-binding site. We present high-resolution structures of *Deinococcus radiodurans* Nramp in three stable conformations of the transport cycle revealing that global conformational changes are supported by distinct coordination geometries of its physiological substrate, Mn^2+^, across conformations and conserved networks of polar residues lining the inner and outer gates. A Cd^2+^-bound structure highlights differences in coordination geometry for Mn^2+^ and Cd^2+^. Measurements of metal binding using isothermal titration calorimetry indicate that the thermodynamic landscape for binding and transporting physiological metals like Mn^2+^ is different and more robust to perturbation than for transporting the toxic Cd^2+^ metal.

## Introduction

Transition metal ions such as Mn^2+^ and Fe^2+^ are essential for various metabolic processes in all living cells and are usually required in low intracellular concentrations for optimal activity^1, 2^. Excess or deficiency of transition metal ions leads to diseases^3, 4^. For example, Fe^2+^ deficiency causes anemia and neurodegenerative diseases whereas Fe^2+^ overload increases the risk of cancer by generating toxic reactive oxygen species (ROS) and mutations^5, 6^. Mn^2+^ overload in the brain is linked to neurological disorders and deficiency causes metabolic defects and impairs growth^7, 8^. Other transitions metals, like Cd^2+^ and Hg^2+^, are toxic and their accumulation affects health by disrupting the physiological levels of essential metals or displacing them in enzyme active sites, thus inhibiting the proteins, or changing their activity^1, 9^. Cells and organisms have evolved strategies to maintain metal ion homeostasis via highly regulated transport and storage processes^3, 4, 10^.

Natural resistance-associated macrophage proteins (Nramps) are ubiquitous importers of Fe^2+^ and Mn^2+^ across cellular membranes into the cytosol^2, 10, 11^. In humans, Nramp1 extrudes essential metals from phagosomes of macrophages to aid in killing engulfed pathogens, and Nramp2 (DMT1) is expressed at low levels in the endosomes of all nucleated cells and imports Mn^2+^ and Fe^2+^ into the cytosol^12–14^. Plant and fungal Nramps aid in Fe^2+^ and Mn^2+^ uptake and trafficking, and bacterial Nramps are involved in the acquisition of Mn^2+^, an essential nutrient^2^. In addition to the physiological substrates Fe^2+^ and Mn^2+^, Nramps can also transport toxic metals like Cd^2+^ and Hg^2+^, but exclude the abundant alkaline earth metals like Mg^2+^ and Ca^2+^ (ref ^2^).

Recent bacterial Nramp structures reveal a LeuT fold, three stable conformations (outward-open, occluded, and inward-open), and identify the metal-binding site residues, including conserved aspartate, asparagine, and methionine residues^15–18^. The metal-binding methionine is essential to select against alkaline earth metals^19^. This finding is corroborated by the fact that a bacterial Nramp homolog which lacks a metal-binding methionine, NRMT, can transport Mg^2+^ (ref ^20^). However, little is known about whether from their promiscuous spectrum of transition metal substrates, the canonical Nramps can mechanistically distinguish between their physiological substrates (Fe^2+^ and Mn^2+^) from non-essential ones like Cd^2+^. Functional studies on *Deinococcus radiodurans* (Dra)Nramp revealed that Mn^2+^ and Cd^2+^ transport show differences in their dependence on pH, proton flux, and membrane potential^17, 21^. However, we the lack of high-resolution structural information on binding of different metals to explain these differences.

We present high-resolution structures of DraNramp in three conformations in both Mn^2+^-bound and metal-free states, providing the first molecular map of the entire Mn^2+^ transport cycle. The structures along with molecular simulations reveal distinct Mn^2+^-coordination spheres and key polar-residue networks that gate the inner and outer vestibules to achieve alternate access during transport. This Mn^2+^ transport cycle also informs on the transport of Fe^2+^, the other common physiological Nramp substrate, because Fe^2+^ and Mn^2+^ have similar coordination preferences and chemical properties^2, 22–24^. Comparisons with an additional high-resolution structure of DraNramp bound to a non-physiological substrate, Cd^2+^, and complementary binding and transport measurements and mutational analyses, suggest that Nramps can distinguish physiological from toxic substrates through thermodynamic differences in the conformational landscape of the transport cycle.

## Results

### DraNramp transports a mostly dehydrated Mn^2+^ ion

To visualize how the metal substrate is coordinated in Nramp transporters, we determined several high-resolution crystal structures of DraNramp using lipid-mesophase based techniques (Supplementary Table 1). We obtained a structure of wildtype (WT) DraNramp in an occluded state bound to Mn^2+^ at 2.38 Å by soaking crystals with Mn^2+^ (WT•Mn^2+^; Table 1, Fig. 1a-b). We resolved a comparable structure at 2.39 Å using the inward-locking mutation A47W^18^ and co- crystallization with Mn^2+^ (A47W•Mn^2+^; Supplementary Table 2; Cα RMSD = 0.47 Å; all pairwise RMSD values listed in Supplementary Table 3). The similarity of both structures, including a nearly identical Mn^2+^ coordination sphere (Fig. 1b, Extended Data Fig. 1a), suggest that the observations we make based on these two structures are robust. Both structures superimpose best with the previously determined occluded metal-free G45R structure^17^ (Cα RMSD of 0.57 and 0.42 Å for WT•Mn^2+^ and A47W•Mn^2+^ respectively); we generally used the WT•Mn^2+^ structure for analysis of the occluded state. As in the metal-free G45R^17^, the Mn^2+^- bound occluded structures have a completely sealed outer vestibule and a partially closed inner vestibule, with the Mn^2+^ occluded from bulk solvent (Fig. 1a).

**Figure 1.**
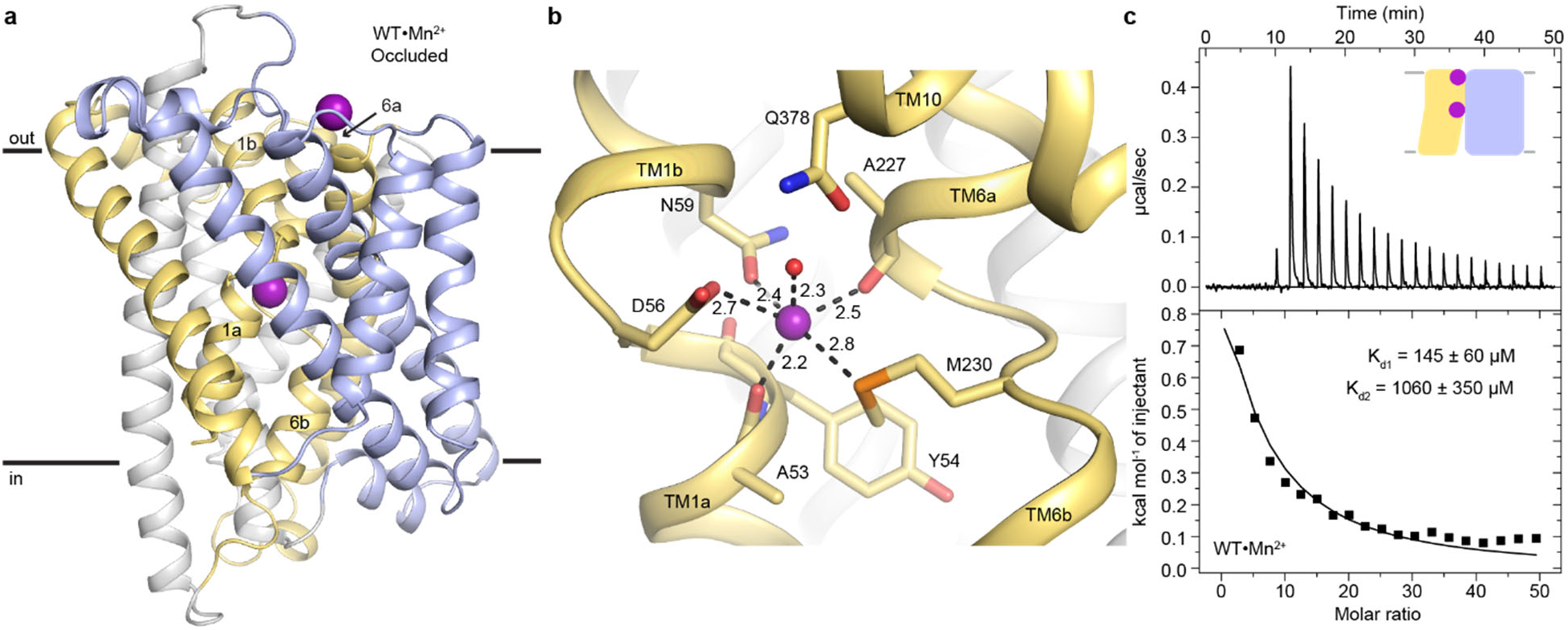
Occluded structure of DraNramp reveals a largely dehydrated Mn^2+^ coordination sphere. (a) Cartoon representation of WT•Mn^2+^ in an occluded state. Anomalous signal confirmed the presence of Mn^2+^ in both the orthosteric metal-binding site and an additional site at the mouth of the external vestibule which is less conserved across the Nramp family (Supplementary Figure 2). TM1 and TM6 are labeled. (b) Detail of the orthosteric metal- binding site of WT•Mn^2+^ where D56, N59, M230, and the pseudo-symmetrically related carbonyls of A53 and A227 coordinate the Mn^2+^ ion. A water molecule completes the 6-ligand coordination sphere. Coordinating residues are shown as sticks, and coordinating distances are indicated in Å. (c) ITC measurement of the affinity of WT DraNramp for Mn^2+^. Top graph shows heat absorbed upon injection of Mn^2+^ solution to the protein solution. Bottom graph shows the fit of the integrated and corrected heat to a binding isotherm. The data show an endothermic mode of binding and were fit using a two-binding-site model (K_d1_ = 145 ± 60 µM, K_d2_ = 1060 ± 350 µM). Based on ITC experiments comparing Mn^2+^ binding to WT or DraNramp constructs with mutations at the external site (Extended Data Fig. 1), we assigned K_d1_ to the orthosteric site. In all figures, unless otherwise noted, TMs 1, 5, 6 and 10 are pale yellow, TMs 2, 7 and 11 gray, TMs 3, 4, 8, and 9 light blue, and Mn^2+^ atoms are magenta spheres.

**Table 1.** Data collection and refinement statistics for four new DraNramp structures

| Structure | WT<sub>soak</sub> | WT•Mn<sup>2+</sup> | M230A•Mn<sup>2+</sup> | WT•Cd<sup>2+</sup> |
| --- | --- | --- | --- | --- |
| Conformation | Occluded | Occluded | Inward open | Inward open |
| Bound metal ion substrate | none | Mn<sup>2+</sup> | Mn<sup>2+</sup> | Cd<sup>2+</sup> |
| PDB ID | 8E5V | 8E60 | 8E6I | 8E6M |
| **Data Collection** |  |  |  |  |
| Beamline | GMCA 23IDB | GMCA 23IDB | GMCA 23IDB | NECAT 24IDC |
| Wavelength (Å) | 1.033 | 1.033 | 1.033 | 0.984 |
| Resolution range (Å) | 41.23-2.36 (2.44-2.36) | 41.28-2.38 (2.46 - 2.38) | 45.32-2.52 (2.61-2.52) | 45.54-2.48 (2.57-2.48) |
| Space group | P 2 21 21 | P 2 21 21 | P 2 21 21 | P 2 21 21 |
| Unit cell (*a*, *b*, *c*) | 58.95, 71.04, 98.77 | 59.08, 71.10, 98.75 | 58.67, 71.35, 98.59 | 59.14, 71.37, 99.05 |
| Unit cell ($\alpha$, $\beta$, $\gamma$) | 90, 90, 90 | 90, 90, 90 | 90, 90, 90 | 90, 90, 90 |
| Number of crystals | 1 | 1 | 3 | 1 |
| Total reflections | 58744 (5928) | 57472 (5733) | 146077 (14913) | 76829 (7275) |
| Unique reflections | 17477 (1718) | 16468 (1646) | 14548 (1427) | 15351 (1507) |
| Redundancy | 3.4 (3.4) | 3.5 (3.5) | 10.0 (10.4) | 5.0 (4.8) |
| Completeness (%) | 98.71 (98.85) | 95.11 (96.92) | 99.90 (99.79) | 99.03 (99.47) |
| Mean *I*/$\sigma$(*I*) | 8.89 (0.97) | 8.92 (0.89) | 8.47 (0.75) | 9.90 (1.12) |
| Wilson *B*-factor | 49.97 | 50.55 | 54.29 | 49.52 |
| *R*<sub>merge</sub> | 0.106 (1.292) | 0.109 (1.241) | 0.269 (2.475) | 0.158 (1.626) |
| *R*<sub>meas</sub> | 0.127 (1.511) | 0.127 (1.447) | 0.284 (2.603) | 0.178 (1.831) |
| *R*<sub>pim</sub> | 0.067 (0.759) | 0.063 (0.722) | 0.090 (0.800) | 0.078 (0.816) |
| CC 1/2 | 0.99 (0.37) | 0.99 (0.39) | 0.98 (0.34) | 0.99 (0.34) |
| **Refinement** |  |  |  |  |
| Resolution range (Å) | 41.23-2.36 (2.44-2.36) | 41.28-2.38 (2.46 - 2.38) | 45.32-2.52 (2.61-2.52) | 45.54-2.48 (2.57-2.48) |
| No. reflections | 17441 (1714) | 16438 (1636) | 14547 (1425) | 15291 (1507) |
| No. reflections in *R*<sub>free</sub> | 1743 (171) | 1642 (164) | 1454 (143) | 1530 (151) |
| *R*<sub>work</sub> | 0.217 (0.340) | 0.207 (0.316) | 0.225 (0.313) | 0.202 (0.319) |
| *R*<sub>free</sub> | 0.245 (0.350) | 0.259 (0.358) | 0.266 (0.349) | 0.250 (0.354) |
| Number of atoms | 3449 | 3385 | 3451 | 3321 |
| Protein | 2945 | 2933 | 2934 | 2905 |
| Ligand | 443 | 405 | 448 | 362 |
| Water | 61 | 47 | 69 | 54 |
| Protein Residues | 392 | 393 | 392 | 388 |
| Ramachandran plot |  |  |  |  |
| Favored (%) | 98.46 | 98.47 | 98.21 | 98.96 |
| Allowed (%) | 1.54 | 1.53 | 1.79 | 1.04 |
| Outliers (%) | 0 | 0 | 0 | 0 |
| Rotamer outliers (%) | 0.33 | 1.00 | 1.01 | 1.01 |
| Clashscore | 8.25 | 8.97 | 7.15 | 5.57 |
| RMS (bonds) | 0.002 | 0.002 | 0.002 | 0.002 |
| RMS (angles) | 0.43 | 0.46 | 0.43 | 0.46 |
| Average *B*-factor | 65.12 | 64.98 | 66.61 | 64.82 |
| Protein | 63.36 | 63.23 | 65.04 | 62.68 |
| Ligand | 77.99 | 78.48 | 77.70 | 83.11 |
| Water | 56.34 | 58.24 | 61.33 | 57.45 |
| No. of TLS groups | 9 | 8 | 3 | 3 |

Anomalous difference Fourier maps confirmed presence of Mn^2+^ at the canonical, orthosteric Nramp metal-binding site between the unwound regions of TM1 and TM6 (Supplementary Table 4, Extended Data Fig. 1a). In WT•Mn^2+^, the Mn^2+^ is coordinated by conserved D56 (2.7 Å), N59 (2.4 Å), M230 (2.8 Å) and backbone carbonyls of A53 (2.2 Å) and A227 (2.5 Å; Fig. 1b, Supplementary Table 5). A53 and A227 are pseudosymmetrically related in the inverted repeats of the LeuT fold of DraNramp (Extended Data Fig. 1b). A water (2.3 Å) bridging Mn^2+^ with Q378, a residue previously proposed to directly coordinate Mn^2+^ in the occluded state^15, 17^, completes the coordination sphere. This yields a coordination number of 6, typical for Mn^2+^, and a largely dehydrated metal-binding site with a distorted octahedral Mn^2+^-coordination geometry (RMS_angles_ = 25°, Supplementary Table 6), as often observed in Mn^2+^-protein complexes^25–27^.

We observed an additional bound Mn^2+^ bridging D296 and D369 at the N-termini of extracellular helix 2 (EH2) and TM10, respectively, at the mouth of the outer vestibule (Extended Data Fig. 1a). We denote this site as the ‘external site’ and the canonical substrate-binding site as the ‘orthosteric site’. Corroborating the structural findings, isothermal titration calorimetry (ITC) measurements reveal an endothermic mode of binding with two Mn^2+^-binding sites for WT (K_d1_ = 145 ± 60 µM, K_d2_ = 1060 ± 350 µM; Fig. 1c) and A47W (K_d1_ = 115 ± 10 µM, K_d2_ = 1355 ± 460 µM; Extended Data Fig. 1d; all K_d_ values are in Supplementary Table 7). To determine the affinity of the orthosteric site, we mutated the external-site aspartates. The D296A and D369A substitutions have little impact on Mn^2+^ transport (Extended Data Fig. 1c). The D296A and D369A variants each bind one Mn^2+^ ion with K_d_ = 240 ± 20 µM and 270 ± 30 µM respectively (Extended Data Fig. 1d). Hence, the affinity for Mn^2+^ at the orthosteric site is ∼5-10 fold higher compared to the external site. A metal ion is present at the external site in all inward-open and occluded metal-bound DraNramp structures, but not outward-open structures, as opening the outer vestibule separates D296 and D369 and disrupts the site (Supplementary Fig. 2a). D296 and D369 are not conserved across Nramps, but they are more conserved within bacterial clade A, and there is a general abundance of acidic residues in the corresponding loop regions across all clades (Supplementary Fig. 2b, c). At present, our results provide little evidence of a biological role for this previously unidentified external site; perhaps the concentration of acidic residues at the mouth of the outer vestibule (Supplementary Fig. 2d) provides electrostatic attraction for metal cations.

### Snapshots of the complete Mn^2+^ transport cycle by DraNramp

We also determined DraNramp structures in metal-free occluded (WT; 2.36 Å) and Mn^2+^-bound inward-open states (M230A•Mn^2+^; 2.52 Å; Table 1, Supplementary Tables 1-2). Along with our previous DraNramp structures^17, 18^, these new structures allow us to map the entire Mn^2+^ transport cycle to three major conformations, each in Mn^2+^-bound and metal-free states (Fig. 2a,b). By ordering and comparing these six structures, we outline a molecular mechanism by which metal substrate binds, induces conformational change, and is released.

**Figure 2.**
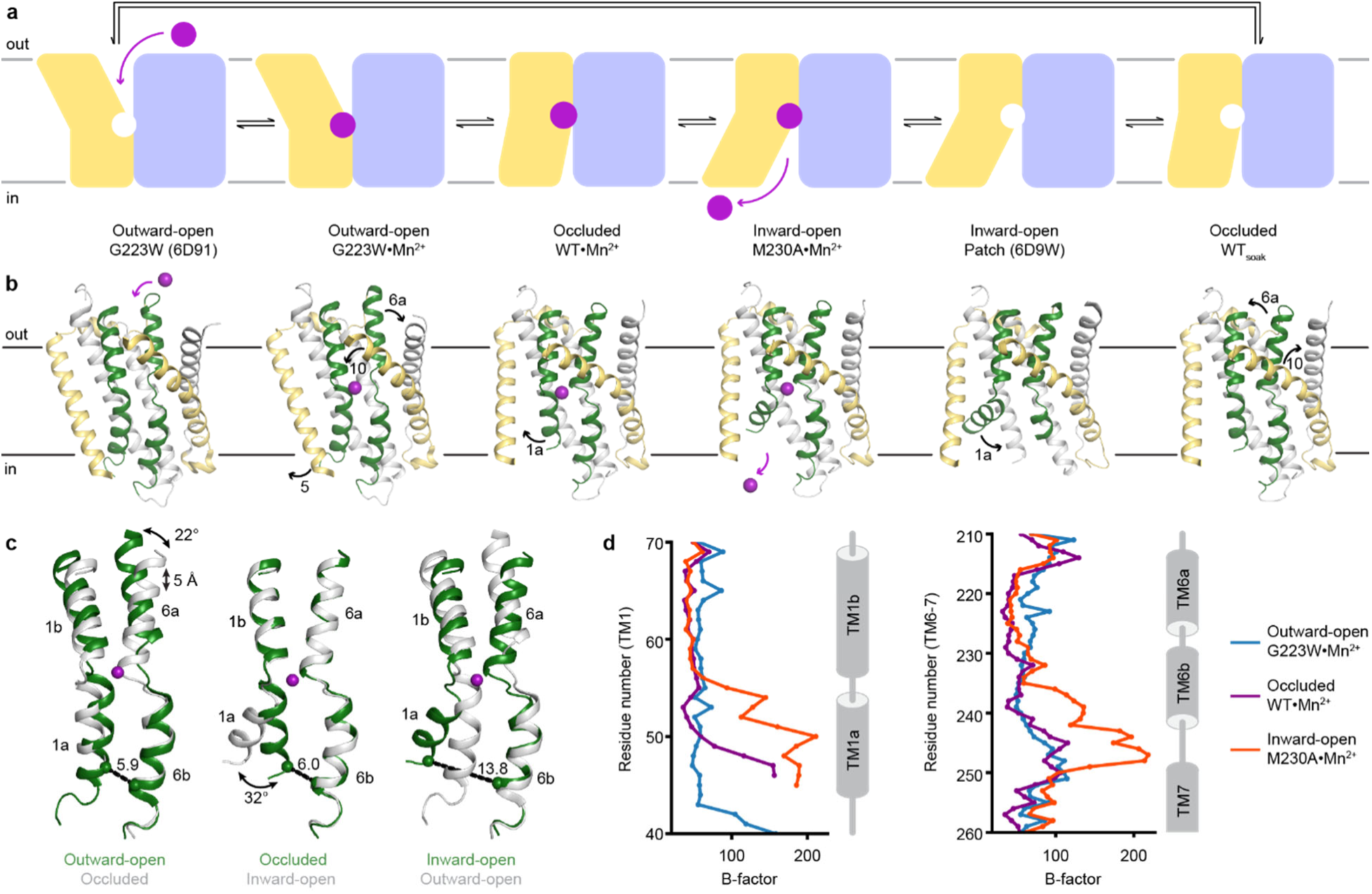
Structures in new conformations complete the Mn^2+^ transport cycle by DraNramp. (a) Schematic of the conformational states that DraNramp traverses to import Mn^2+^. The mobile and stationary parts are pale yellow and light blue, respectively. (b) Corresponding structures of DraNramp, showing TMs 1 and 6 in green, TMs 5 and 10 in pale yellow, and TMs 2, 7 and 11 gray. Stationary TMs 3, 4, 8, and 9 are omitted to highlight the key motions in the mobile parts. Mn^2+^ ions are magenta. Black arrows indicate the key motions in TMs 1a, 5, 6a and 10. Full structures and the electron density for TMs 1 and 6 are illustrated in Extended Data Figure 2. (c) Pairwise superpositions of whole Mn^2+^-bound structures highlight the motions of TMs 1 and 6. Conformations are indicated at the bottom. The distance between residues 46 and 240 in TMs 1a and 6b, indicated for the green structures, increases from 5.9 Å to 13.8 Å from outward open to inward open. The large angular motions of TM1a and TM6b are also indicated. (d) Plots of B-factor by residue for the TM1 region (residues 40-70) and the TM6 region. The B-factor are highest for the inward-open state in which the interaction between TMs 1a and 6b (both in the inner leaflet) is broken.

TMs 1, 5, 6, and 10 move the most as the Mn^2+^-binding site accessibility switches from outward to inward across the conformations (Fig. 2b). As Mn^2+^ binds to the outward-open state, TM10 moves towards TM1b, TM5 towards TM7, and TM6a towards TM11 to seal the outer vestibule and yield an occluded state. The TM5 position in the occluded state is primed for TM1a to swing upward to open the inner vestibule. This swing of TM1a is the only significant difference between the Mn^2+^-bound occluded and inward-open states, allowing release of the Mn^2+^ into the inner vestibule. TM1a swings to a similar angle in the new inward-open M230A•Mn^2+^ as in the previous low-resolution inward-open metal-free structure^18^, and the homologous LeuT and serotonin transporter structures^28, 29^, for example. Thus, most of the structural reorganization in Nramps occurs in the shift from outward open to occluded. The three metal-free DraNramp conformations are similar to their corresponding Mn^2+^-bound structures, suggesting that once Mn^2+^ is released, the conformational transitions are reversed, including passing through an occluded metal-free intermediate, to reach the outward-open metal-free conformation ready to accept Mn^2+^ (Fig. 2a-b, Extended Data Fig. 2a).

The substrate-binding TM1 and TM6 are well-resolved in our structures (Extended Data Fig. 2b) and pairwise superpositions reveal how their motions contribute to the conformational changes across the transport cycle (Fig. 2c). TM6b tilts by 22° and the unwound region of TM6 becomes more helical as it moves towards the Mn^2+^ to close the outer gate. The central unwound regions of TM1 and TM6 are closest in the occluded state, resulting in an almost dehydrated Mn^2+^-coordination sphere. Finally, the inner vestibule opens when TM1a tilts upward by 32°, increasing the distance between TM1a and TM6b by ∼8 Å (Fig. 2c). TM6b is largely static relative to the protein core, although in the inward-open structure it has high B-factors (Fig. 2d), indicating that the interaction with TM1a stabilizes TM6b to close the inner vestibule.

### Different conformations have distinct Mn^2+^ coordinations

Our Nramp structures provide snapshots of the complete Mn^2+^-coordination sphere geometries in each conformation (Fig. 3). Two new structures reveal the coordination of Mn^2+^ in the inward-open state: M230A•Mn^2+^ and D296A•Mn^2+^ (Cα RMSD of 0.42 Å). The Mn^2+^ is in the same location of the orthosteric site as in the occluded state, as confirmed by anomalous diffraction for D296A•Mn^2+^ (Supplementary Table 4, Extended Fig. 3a-b). As in the occluded state, in M230A•Mn^2+^ the Mn^2+^ binds D56 (2.7 Å), N59 (2.6 Å), and the A227 carbonyl (2.4 Å), with a water replacing M230 (2.3 Å; Fig. 3a, Supplementary Table 5). However, with TM1a displaced, the A53 carbonyl no longer coordinates Mn^2+^; instead, the Y54 carbonyl distantly coordinates Mn^2+^ at 3.1 Å. Two more waters, one bound to Q378 (2.5 Å) and another from the inner vestibule (2.2 Å), complete a seven-coordination sphere resembling a pentagonal bipyramidal geometry with significant distortion (RMS_angle_ = 36°, Supplementary Table 6). Seven coordination is infrequent but found in Mn^2+^-coordinating proteins like MntR^30, 31^. Our inward-open structures provide the first evidence that Y54 participates in the Mn^2+^ transport cycle.

**Figure 3.**
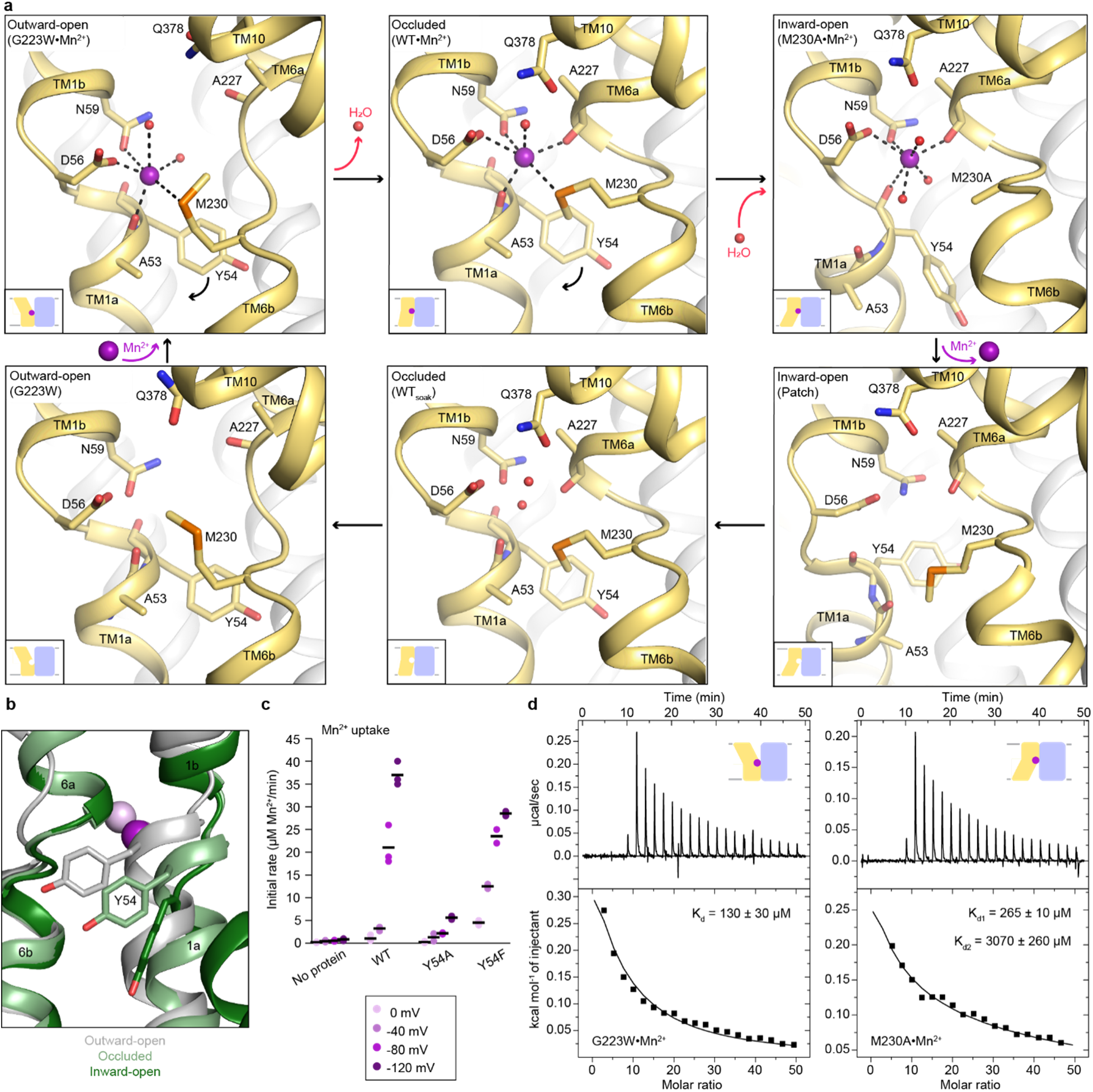
Coordination sphere changes across the Mn^2+^ transport cycle of DraNramp. (a) Structures of the orthosteric metal-binding site in six conformations reveal the differences in coordination geometry and illustrate that the bound Mn^2+^ is more hydrated in the outward-open and inward-open states than the occluded state. In the occluded structure of metal-free WT DraNramp a density we have assigned as water replaces Mn^2+^. Y54 in TM1a progressively moves to open the inner vestibule in the transition from outward to inward open, shown by black curved arrows. (b) TM1 and TM6 from a superposition of the three Mn^2+^-bound structures in panel a illustrate the swing of the Y54 sidechain as sticks. The view is rotated 180° along the vertical axis from Figure 2c. (c) Initial Mn^2+^ uptake rates for DraNramp variants Y54A and Y54F at membrane potentials ranging from ΔΨ = 0 to −120 mV (n = 2-3; black bars are the mean values). The Mn^2+^ concentration was 750 μM, and the pH was 7 on both sides of the membrane. Y54A nearly abolishes transport whereas Y54F has near-wildtype initial transport rates. Corresponding time traces are plotted in Supplementary Figure 1. (d) ITC measurements of the affinity of G223W (left) and M230A (right) for Mn^2+^. The external Mn^2+^-binding residues drift apart in the outward-open structure (Supplementary Fig. 2a), hence, we assign the single Mn^2+^ binding event to the orthosteric site for outward-locked G223W. M230A data fit a two- binding-site model, consistent with its structure and WT DraNramp.

The new occluded and inward-open Mn^2+^-bound structures have a monodentate coordination of D56 with Mn^2+^. For consistency, we reinterpreted the outward-open G223W•Mn^2+^ map (PDB ID: 6BU5) with a monodentate coordination of D56 with Mn^2+^ instead of previously modeled bidentate interaction^17^; the local and global model statistics are very similar to the original structure (Supplementary Table 5). Re-refined G223W•Mn^2+^ has six Mn^2+^-coordinating ligands: D56 (2.4 Å), N59 (3.2 Å), M230 (3.0 Å), carbonyl of A53 (2.4 Å) and two waters (2.7 Å and 2.6 Å; Fig. 3a, Supplementary Table 5); the overall geometry resembles a distorted octahedron as in the occluded structure (RMS_angle_ = 22°; Supplementary Table 6).

Comparing the three Mn^2+^-bound conformations (Fig. 3a), the carbonyls of A53 in TM1a and A227 in TM6b alternately coordinate Mn^2+^ in the outward and inward-open structures respectively, and both residues interact with Mn^2+^ in the occluded structure. The pseudosymmetrically related A53 and A227 may thus act as hinges altering the Mn^2+^- coordination sphere as TM1 and TM6 move in turn to open the gates during Mn^2+^ transport. Furthermore, as DraNramp switches from outward- to inward-open, Y54 progressively swings downward, acting as a gate in concert with TM1a’s upward swing to open the inner vestibule and allow metal release (Fig. 3a-b). Our inward-open structures also suggest that the Y54 carbonyl may participate in Mn^2+^ release through a direct interaction with the metal ion (Fig. 3a). All 3796 Nramp sequences in our alignment have either a tyrosine or a phenylalanine at this position and Y54 is completely conserved among bacterial clades A (including DraNramp) and C, while the position is 40% and 100% phenylalanine among eukaryotes and bacterial clade B, respectively (Extended Data Fig. 3c). To evaluate the significance of Y54 in Mn^2+^ transport, we purified and reconstituted into proteoliposomes the Y54A and Y54F variants. While Y54F has near-wildtype Mn^2+^-transport activity, Y54A nearly eliminates Mn^2+^ transport (Fig. 3c), indicating that an aromatic ring is essential for the gating motion required for transport.

We also measured the Mn^2+^-binding affinity of the constructs that yielded inward- or outward- open structures, M230A and G223W, respectively. Mn^2+^ is present at the external site in the inward-open M230A•Mn^2+^ (Supplementary Fig. 2a), and consistently, the ITC data indicate two Mn^2+^-binding sites in M230A (K_d1_ = 265 ± 10 µM assigned to the orthosteric site, K_d2_ = 3070 ± 260 µM for the external site; Fig. 3d). The reduced affinity of M230A compared to WT is consistent with the absence of the Mn^2+^-coordinating sulfur ligand from M230. The ITC data for G223W indicate a single Mn^2+^-binding site (K_d_ = 130 ± 30 µM; Fig. 3d), which we assign to the orthosteric site because the opening of the outer vestibule displaces TM10, disrupting the external site (Supplementary Fig. 2a). The affinity of G223W for Mn^2+^ is similar to that of WT at the orthosteric site, suggesting that the outward-open and occluded states have similar affinity.

### Mn^2+^ binding does not significantly alter the three main DraNramp conformations

To compare the metal-bound states to analogous metal-free states of the transport cycle, we determined two metal-free occluded structures of wildtype DraNramp at a higher resolution than the previously reported G45R structure^17^, which we refer to as WT (2.38 Å; Supplementary Table 2) and WT_soak_ (2.36 Å; Table 1). The crystal used for WT_soak_ was mock-soaked (with no metal in the soaking solution). WT and WT_soak_ are nearly identical (Cα RMSD of 0.38 Å) confirming that the soaking process does not influence the conformational state. We analyzed WT_soak_, unless otherwise noted. In WT_soak_, we observed density but no anomalous signal at the orthosteric site and modeled a water molecule at the position where Mn^2+^ sits in the occluded state (Fig. 3a). WT_soak_ is nearly identical to WT•Mn^2+^ (Cα RMSD of 0.20 Å), indicating that the occluded conformation is unchanged by the presence of metal ion substrate, and the metal-binding site is instead filled by ordered water molecules.

We used previously reported inward-open (which we refer to as ‘Patch’ because it has a patch of intracellular loop mutations; PDB ID: 6D9W) and outward-open (G223W; PDB ID: 6D91) metal-free structures for analysis of the Mn^2+^ transport cycle (Fig. 3a)^17, 18^. These structures, resolved at lower resolution than the ones described here, have no density at the orthosteric site. This is consistent with a more flexible organization of a metal-binding site open to bulk aqueous solvent. In contrast, metal-free WT has a more ordered orthosteric site, suggesting a stable occluded intermediate in the switch from inward- to outward-open.

### Polar networks latch the gates to achieve alternating access

Vestibules providing access to the orthosteric site from the extracellular or intracellular side alternately open from the motions of DraNramp’s TMs 1, 5, 6 and 10, which form the outer and inner gates during Mn^2+^ transport^17, 32^. To pinpoint protein features that enable these motions, we used our Mn^2+^-bound structures to identify interaction networks with the following attributes: (i) they contain conserved polar residues from at least one of the four mobile helices; (ii) they line the gates; and (iii) they rearrange between the three resolved protein conformations (Fig. 4a).

**Figure 4.**
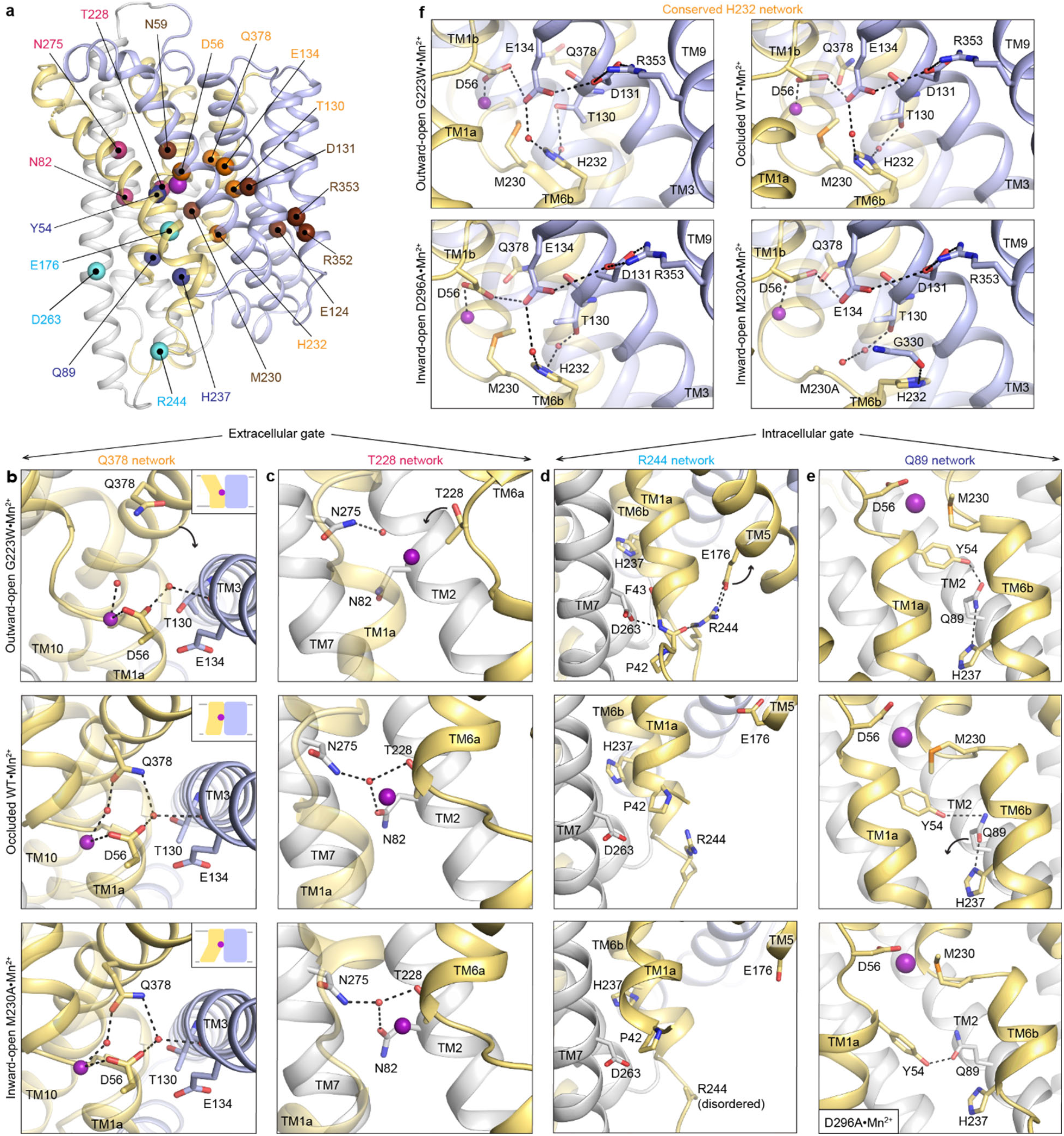
Networks of polar residues lining the outer and inner vestibules rearrange through the conformational transitions needed for Mn^2+^ transport. (a) The Cα positions of residues in the Q378 (orange) and T228 (pink) networks lining the outer gate, R244 (cyan) and Q89 (blue) networks within the inner gate, and the H232 (orange) network coordinating with the proton pathway, are mapped on the occluded WT•Mn^2+^ structure. Metal-binding and proton pathway residues^17^ are represented as brown spheres. (b) The Q378 network forms as TM10 moves when DraNramp transitions from outward-open to occluded to close the outer vestibule. Water-mediated interactions form between Q378, D56, and T130. The other D56 carbonyl interacts directly with Mn^2+^. TM1 is transparent. (c) The T228 network forms with N275, N82, and T228 coordinating a water as TM6a moves to close the outer vestibule. TM1 is transparent. (d) In the R244 network, interactions between R244, E176, and D263 break as TM5 moves in the transition from outward-open to occluded state to initiate the opening of the inner vestibule. (e) In the Q89 network, Y54, Q89, and H237 rearrange from occluded to inward-open state as TM1a swings up to allow for metal release. (f) H232, which abuts the orthosteric Mn^2+^-binding site, interacts with E134 and T130 through waters conserved in all conformations. In the M230A•Mn^2+^ structure, H232 flips and is replaced by a water, retaining the interaction with T130 but breaking the connection with E134, suggesting that M230 helps stably position H232. In panels c-e, TMs 3, 4, 8, and 9 are omitted to better visualize the interactions. The illustrated structures are indicated on the figure.

Two networks seal the outer gate. In the occluded and inward-open conformations, Q378 (TM10) interacts with two waters, one coordinating the orthosteric Mn^2+^ and the other interacting with D56 (TM1) and the carbonyl of T130 (TM3). This network is disrupted in the outward-open conformation as Q378 and the rest of TM10 swing outward to open the outer vestibule (Fig 4b). In the second network, T228 (TM6a), N275 (TM7) and N82 (TM2) interact via a water in the occluded and inward-open states, but not in the outward-open state, where the extended unwound region of TM6a positions T228 farther from the orthosteric site and N275 and N82 (Fig 4c). This T228 network helps rearrange TM6a, closing the outer vestibule in the occluded state and generating a nearly dehydrated Mn^2+^-coordination sphere (Fig. 1b). As the inner gate opens to release Mn^2+^, both networks persist, ensuring that the outer gate remains closed in the inward-open conformation.

All six residues in the Q378 and T228 networks are completely conserved across bacterial clade A and highly conserved across all Nramps (Supplementary Fig. 3). To investigate the robustness of these networks, we performed duplicates of molecular dynamics (MD) simulations starting in all three conformations and confirmed that within the first 250 ns of these simulations, the T228 and Q378 networks persist in simulations of the occluded and inward-open states and remain broken in simulations starting in the outward-open state (Extended Data Fig. 4a-c). This includes the coordinated waters, for example the water at the center of the Q378 network is present in more than 50% of the frames in occluded-state simulations (Extended Data Fig 4e). In line with the X-ray snapshots and simulation outcomes, mutations of any of the six residues across these networks reduces Mn^2+^ transport by DraNramp^17, 19, 21, 32^, highlighting their key function in the conformational cycle of Nramps.

The inner vestibule is gated by rearrangements of residues in two other polar networks, namely those of R244 and Q89 (Fig. 4d-e). In the outward-open state, R244 (TM6b) forms an ion pair with E176 (TM5) and interacts with the TM1a backbone, as does D263 (TM7), keeping TM1a and TM5 close and the inner gate closed (Fig. 4d)^32^. In the occluded state, the E176-R244 interaction breaks and TM5 moves away from TM6b, creating space for TM1a to swing up and open the inner vestibule in the inward-open state. Accordingly, the E176–R244 ion pair is stable in MD simulations of the outward-open state, while these residues are >10 Å apart in simulations of the occluded and inward-open states (Extended Data Fig. 4d). Supporting the importance of the R244 network, E176 is 100% and R244 is 85% conserved across all Nramps (Supplementary Fig. 3) and mutation of either residue reduces Mn^2+^ transport by DraNramp^32^.

In the Q89 network, Q89 (TM2) hydrogen-bonds with Y54 (TM1a) and H237 (TM6b) to seal the inner gate in the outward-open^32^ and occluded states (Fig. 4e). In the inward-open structure, the Q89–H237 hydrogen bond is broken and both Y54 and Q89 rearranged into a different hydrogen bond, buttressing the opening of the inner vestibule (Fig. 4e). As discussed above, Y54 is conserved and important for Mn^2+^ transport by DraNramp (Fig. 3c and Extended Data Fig 3c).

Similarly, Q89 and H237 are conserved (Supplementary Fig. 3), and mutations to either residue affect Mn^2+^ transport in a cell-based assay^32^. Cysteine accessibility measurements showed that mutations to Q89 or H237 render the outer vestibule solvent-inaccessible^32^, indicating that disrupting the Q89 network likely prevents closing of the inner gate.

H232 (TM6b) is conserved across all Nramps (Supplementary Fig. 3), highlighting its importance. H232 sits below the orthosteric site and forms a network conserved across all conformations, with water-mediated hydrogen bonds to E134 (involved in proton transfer to the salt-bridge residues in TMs 3 and 9)^17, 21^ and T130 in TM3 (Fig. 4f). Waters occupy these two sites in MD simulations in all states, especially the water coordinated between H232 and T130 (Extended Data Fig. 4f-g). However, in M230A•Mn^2+^ H232 flips to interact with the G330 carbonyl (TM8; Fig. 4f), suggesting that it may transiently move during the conformational cycle. Indeed, while the H232 sidechain rotamer is stable in MD simulations of the outward-open state, it explores other rotamers in simulations of the occluded and inward-open states (Supplementary Fig. 4).

### Unlike Mn^2+^ binding, Cd^2+^ binding to DraNramp is exothermic

Nramps transport divalent transition metals quite promiscuously, including both physiological (Fe^2+^ and Mn^2+^) and non-physiological substrates (Cd^2+^, Zn^2+^, Co^2+^, Ni^2+^, Pb^2+^), but select against alkaline earth metals (Mg^2+^, Ca^2+^)^2, 16, 17, 19, 33–35^. DraNramp transports Cd^2+^ well, but without concomitant proton flux and with weaker voltage dependence (Extended Data Fig. 5a)^2, 17, 21^. To better understand the underlying mechanistic differences, we compared the binding, transport, and structures of DraNramp with Mn^2+^ and Cd^2+^.

**Figure 5.**
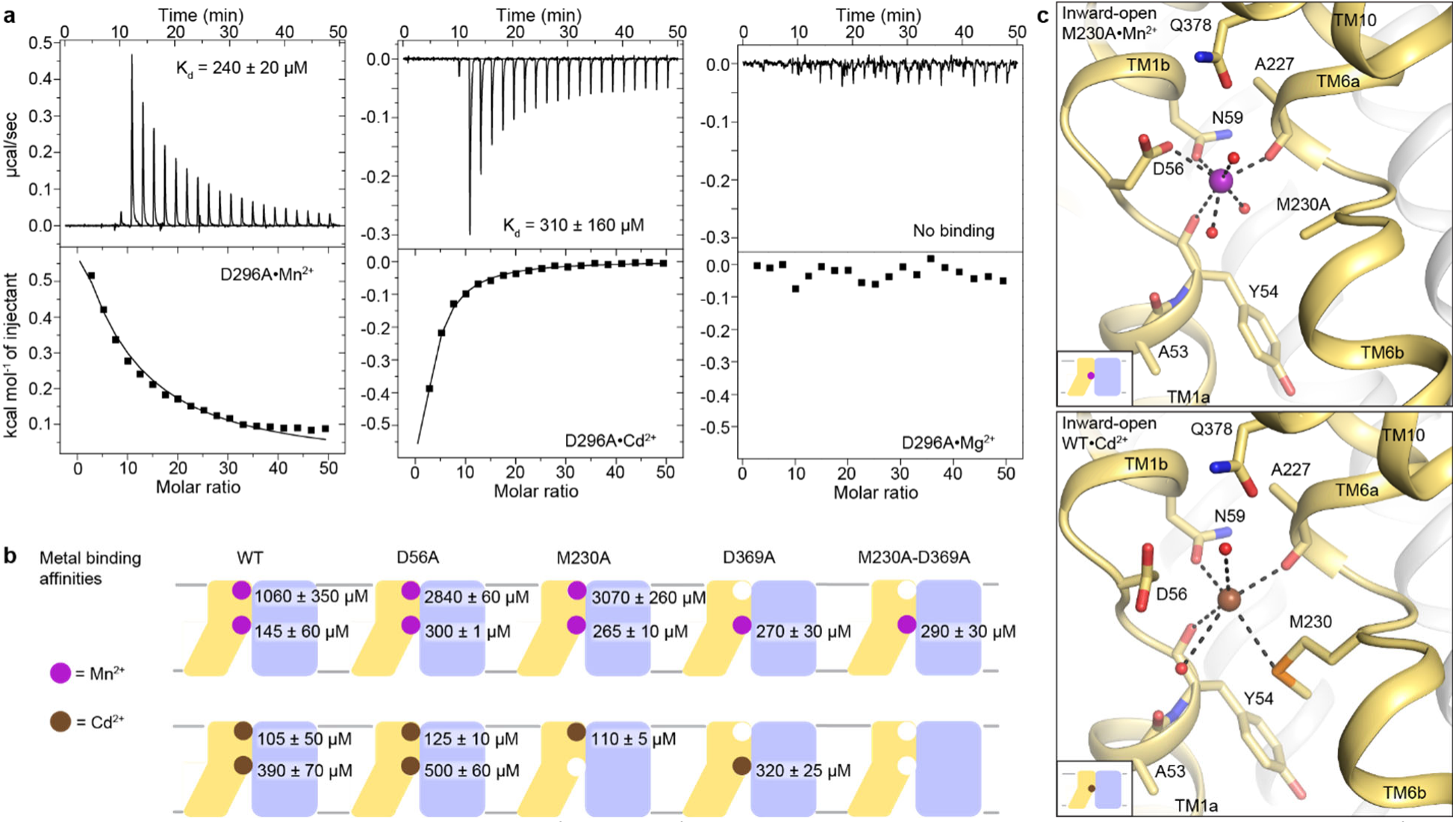
DraNramp binds differently to Mn^2+^ and Cd^2+^. (a) ITC measurements show that DraNramp binds Mn^2+^ in an endothermic mode, Cd^2+^ in an exothermic mode and does not bind Mg^2+^ (6 mM metal). ITC was performed with DraNramp in which the external site is mutated (D296A) to analyze binding at the orthosteric site. (b) Schematic showing the ITC-measured K_d_ values of various DraNramp constructs for Mn^2+^ (top) and Cd^2+^ (bottom). WT DraNramp binds Mn^2+^ and Cd^2+^ at the same two sites (Fig. 1 and Extended Data Fig. 5), with the external-site affinity 10-fold higher for Cd^2+^ than Mn^2+^, whereas the orthosteric-site affinity is higher for Mn^2+^. D56A retains binding at the orthosteric site albeit with weakened affinity for both metals, although D56 coordinates Mn^2+^ but not Cd^2+^. The M230A mutation eliminates binding of Cd^2+^ but not Mn^2+^ at the orthosteric site. The D369A mutation eliminates binding of either metal at the external site. A variant with mutations at both the orthosteric and external sites, M230A-D369A, does not bind Cd^2+^ but maintains orthosteric site binding for Mn^2+^. Additional ITC traces are shown in Extended Data Figure 5b and 5e-f. (c) Comparison of the inward-open state bound to Mn^2+^ (M230A•Mn^2+^; top) and Cd^2+^ (WT•Cd^2+^; bottom) shows differences in coordination geometry at the orthosteric site. D56 is oriented differently and does not directly coordinate Cd^2+^. Cd^2+^ coordinates N59, M230, carbonyls of A227 and Y54, and two waters, for a total of six ligands compared to seven for Mn^2+^. Mn^2+^ and Cd^2+^ are magenta and brown spheres, respectively. Of note, we use M230A•Mn^2+^ here because soaking of WT crystals with Cd^2+^ yielded an inward-open WT•Cd^2+^ structure (Extended Data Fig. 5c) whereas soaking the same kind of crystals with Mn^2+^ yielded an occluded WT•Mn^2+^ structure (Fig. 1).

In contrast to endothermic binding of Mn^2+^, ITC measurements show exothermic binding of Cd^2+^ to WT DraNramp (Extended Data Fig. 5b). Like for Mn^2+^, the Cd^2+^ isotherm fits a two-binding-site model (K_d1_ = 105 ± 50 µM, K_d2_ = 390 ± 70 µM). Both D296A and D369A—containing mutations at the external metal-binding site—showed the similar exothermic trend as the WT construct but fit a single-site model (Fig. 5a, Supplementary Table 7). Based on their affinity (K_d_ = 310 ± 160 µM for D296A and K_d_ = 320 ± 25 µM for D369A), we conclude that the orthosteric site has lower affinity towards Cd^2+^ than the external site (Fig. 5a-b, Supplementary Table 7). The orthosteric site shows ∼2-fold higher affinity towards the physiological substrate Mn^2+^ (K_d_ of 145 ± 60 µM for Mn^2+^ vs. 390 ± 70 µM for Cd^2+^) whereas the external site has 10-fold higher affinity towards Cd^2+^ (K_d_ of 1060 ± 350 µM for Mn^2+^ vs. 105 ± 50 µM for Cd^2+^). Consistent with its inability to transport Mg^2+^, a representative alkaline earth metal^2, 17^, DraNramp does not bind Mg^2+^ (Fig. 5a).

### A Cd^2+^-bound structure helps explain functional differences

To understand the differences in binding and transport, we determined a 2.5-Å Cd^2+^-bound structure by soaking WT DraNramp crystals with 2 mM Cd^2+^ (Table 1). In agreement with the ITC data, WT•Cd^2+^ shows Cd^2+^ ions at both the external and orthosteric sites as confirmed by anomalous signal (Extended Data Fig. 5c-d, Supplementary Table 4). The ∼10-fold higher affinity of the external site for Cd^2+^ than Mn^2+^ suggests that local geometry (accessible coordination distances and angles) favors non-physiological Cd^2+^ over the physiological substrate Mn^2+^ (Supplementary Fig. 2a). The low affinity of this site for Mn^2+^, compared to Cd^2+^, is consistent with the idea that this external site plays a role in general electrostatic attraction of substrate to the orthosteric site, rather than as a finely tuned binding site specific for Mn^2+^.

Interestingly, WT•Cd^2+^ is inward open, although both mocked-soaked and Mn^2+^-soaked crystals under otherwise equivalent conditions yielded occluded structures (WT_soak_ and WT•Mn^2+^, respectively; Figs. 1 and 3). The larger ionic radius of Cd^2+^ (∼0.95 Å) compared to Mn^2+^ (∼0.82 Å)^36, 37^ and different preferred coordination geometry may increase the stability of the inward-open state with Cd^2+^ bound. The ∼2-fold higher affinity of the orthosteric site for Mn^2+^ over Cd^2+^, consistent with DraNramp’s physiological role, suggests a more favorable Mn^2+^ coordination. Comparing with the inward-open M230A•Mn^2+^, D56 adopts a different rotamer in WT•Cd^2+^ and does not coordinate the Cd^2+^ bound at the orthosteric site (Fig. 5c). The rest of the coordination sphere is similar and includes N59 (2.9 Å), M230 (3.4 Å), the A227 carbonyl (2.8 Å), the Y54 carbonyl (3.2 Å), a water (2.9 Å) that coordinates Q378 and another water (3.3 Å) from inner vestibule. Cd^2+^ has six coordinating ligands, and the distortion from ideal octahedral geometry is more pronounced compared to Mn^2+^ (RMS_angle_ = 34°, Supplementary Table 6). The coordination distances are larger for Cd^2+^ than for Mn^2+^ (Supplementary Table 5), as seen in other proteins^38, 39^ and consistent with its larger ionic radius and distinct charge distribution. Cd^2+^ is a soft metal, likely explaining why it retains coordination by the softer sulfur ligand of M230^40^ but not the hard oxygen of D56, although D56 can still provide favorable electrostatics. ITC of M230A corroborates the structural data with Mn^2+^ binding at two and Cd^2+^ at one site (Fig. 5b and Extended Data Fig 5). M230 is thus a crucial ligand for Cd^2+^ but not Mn^2+^, which is further reflected in the transport behavior where M230A affects Cd^2+^ transport drastically but has negligible effect on Mn^2+^ (ref ^19, 21^).

### Cd^2+^-bound DraNramp is most stable in the inward-open state

At the orthosteric site, Mn^2+^ binding is entropy-driven, whereas Cd^2+^ binding is enthalpy-driven, and entropy contributions values (-TΔS) are ∼5-fold smaller for Cd^2+^ than Mn^2+^ (Supplementary Tables 8-9), suggesting that Cd^2+^ binding conformationally constrains the protein more, leads to less solvent release, or both^41, 42^. ITC data analysis of different DraNramp variants highlights additional differences between Mn^2+^ and Cd^2+^ binding (Fig. 5b, Supplementary Table 7). D56A shows two-site binding with both metals, with all affinity values about two-fold worse than WT, although D56A has essentially no transport activity^17^. This suggests that D56 is more important for catalyzing transport than for substrate binding. As expected, variants with an additional mutation at the external site, D56A-D296A and D56A-D369A (and M230A-D296A and M230A-D369A), yielded single-site Mn^2+^-binding isotherms, which we assigned as binding to the orthosteric site. However, all four double mutants showed complete loss of Cd^2+^ binding, which is expected for ones with M230A (which eliminates Cd^2+^ binding at the orthosteric site on its own, as we have also shown previously^19^), but more surprising for ones with D56A. Thus, binding of the preferred physiological Mn^2+^ substrate at the orthosteric site is more robust to perturbations than binding of Cd^2+^, a toxic metal.

The conformation-locking mutants—A47W, which is outward-closed^18^ and crystallized in an occluded state, and outward-open G223W^17^—also show metal-specific binding behavior. ITC with Mn^2+^ agrees with the structural data, with two-site fit for A47W (Extended Data Fig. 1d), and one-site fit for A47W-D296A, A47W-D369A, and G223W (assigned to the orthosteric site; Extended Data Fig. 5e, Supplementary Table 7). However, with Cd^2+^, A47W shows one-site binding and no binding with A47W-D296A, A47W-D369A, and G223W (Extended Data Fig. 5f and Supplementary Table 7), indicating that there is no significant affinity for Cd^2+^ at the orthosteric site of A47W and G223W. This is consistent with our inability to obtain Cd^2+^-bound structures in conformations other than inward-open, despite trying both co-crystallization and soaking with A47W and G223W. These data suggest that Cd^2+^ has highest affinity for the inward-open state and low affinity for other states, in agreement with the fact that soaking of WT crystals (which yielded the metal-free occluded WT_soak_ structure) produced an inward-open Cd^2+^- bound structure.

### Gating network differences in the Cd^2+^-bound structure are restricted to D56

To better understand the Cd^2+^ transport mechanism given that its binding at the orthosteric site seems less optimal and robust than Mn^2+^, we compared the gating networks described above for the inward-open Mn^2+^- and Cd^2+^-bound structures. The Q89, T228, and R244 networks are essentially identical in the inward-open state, regardless of whether Mn^2+^ or Cd^2+^ is bound (Extended Data Fig. 6a-c).

**Figure 6.**
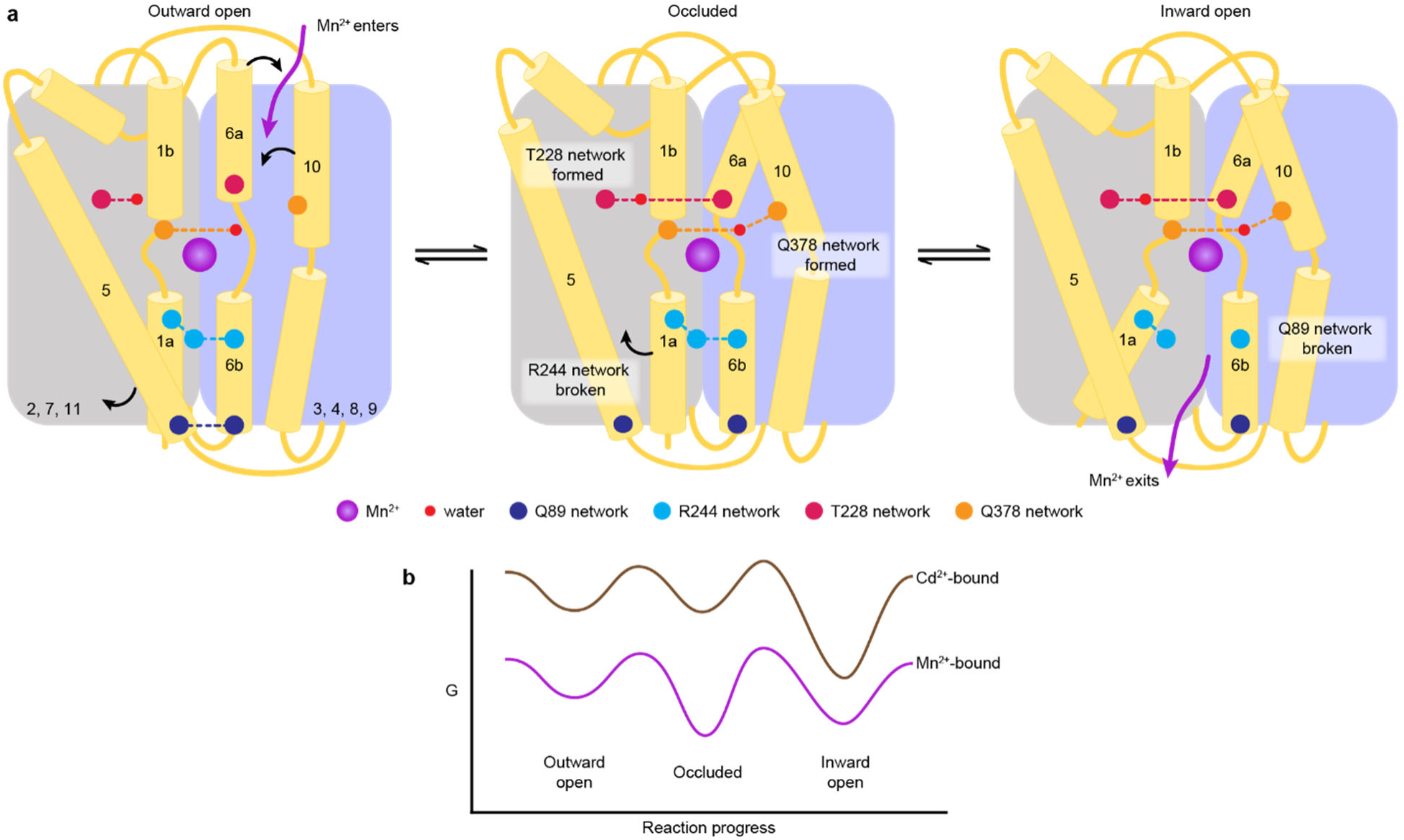
A structure-guided model of the conformational cycle and thermodynamic landscape of metal transport by Nramps. (a) After Mn^2+^ enters through the outer vestibule between TM6a and TM10 in the outward-open state, a bulk conformation change closes the outer gate. The occluded conformation arises though rearrangements of TM6a and TM10 facilitated by formation of the T228 and Q378 networks, respectively. The inner gate partially opens in the occluded state as the R244 network breaks and TM5 moves. To achieve the inward-open conformation, disruption of the Q89 network frees TM1a to swing up to fully open the inner vestibule for Mn^2+^ release into the cytosol. (b) Our data indicate that the most stable Mn^2+^-bound state is the occluded state, and the three main states are readily accessible to facilitate transport. In contrast, Cd^2+^ binding stabilizes the inward-open state.

The other two networks, both of which involve D56, have some differences. In the Q378 network gating the outer vestibule, D56 does not coordinate Cd^2+^ but does interact with the conserved water that connects the metal ion with Q378 in all outward-closed structures (Extended Data Fig. 6d). This preserves a connection to both Cd^2+^ and Q378 to close the outer gate. Most of the H232 network is similar in the Mn^2+^- and Cd^2+^-bound structures, except for the orientation of D56 relative to E134 (Extended Data Fig. 6e).

Mutations of several residues in these polar networks reduced transport in proteoliposome-based assays (Extended Data Fig. 6f), corroborating results from cell-based assays^32^. Y54A, H237A and H232A mutations cause the largest decreases, Q89A causes a moderate decrease, and Y54F is similar to WT. Mn^2+^ and Cd^2+^ follow similar trends although with less voltage dependence for Cd^2+^ than Mn^2+^. These results support the idea that although Mn^2+^ and Cd^2+^ bind differently at the orthosteric site, the flexibility of the D56 sidechain enables the networks of polar residues that gate the outer and inner vestibules to engage and enable transport of both metals at similar rates.

## Discussion

Through detailed analysis of high-resolution structures in different conformations combined with thermodynamic analyses of substrate binding, we provide a molecular map of the Mn^2+^ import pathway in DraNramp. Binding of the Mn^2+^ substrate at the orthosteric site takes on different coordination geometries through the three main conformational states in the transport cycle, all of which deviate significantly from ideal, consistent with the moderate binding affinities we measured. We identified several networks of polar interactions that gate both the outer and inner vestibules. Structures of DraNramp bound to Mn^2+^ and Cd^2+^ and corresponding analyses of substrate binding demonstrate that the orthosteric site shows selectivity for physiological (Mn^2+^) over toxic (Cd^2+^) substrates in binding affinity and robustness to various perturbations.

Our structures of one Nramp homolog, DraNramp, in all conformations of the transport cycle provide an opportunity to update the overview of the conformational cycle, focusing on the polar interaction networks that gate the outer and inner vestibules (Fig. 7a). The structures also provide the first molecular view of how local changes in the coordination spheres and the global conformational transitions coordinate to facilitate Mn^2+^ transport. Starting in the outward-open state, Mn^2+^ entry into the outer vestibule and binding at the orthosteric site triggers the transition to the occluded state by two major rearrangements: (i) closing of the outer gate as the T228 and Q378 networks form and support the reorientation of TM6a and TM10, respectively, and (ii) partial opening of the inner gate through motion of TM5 and breaking of the R244 network. TM6a and TM10 approach the orthosteric site to directly (A227 in TM6a) or indirectly (Q378 in TM10 through a conserved water) coordinate Mn^2+^ and restrict solvent access from the extracellular side in the occluded structure. A227 replaces a water of the outward-open Mn^2+^- coordination sphere, thus retaining a six-coordination geometry in both conformations. Mn^2+^ prefers octahedral (six) coordination^26, 31, 43^ as in the outward-open and occluded structures, although we observed distortions (Supplementary Table 6).

The inward-open state is achieved when TM1a swings up, rupturing the Q89 network and opening the inner gate. The coordination sphere changes again as a water from the open inner vestibule replace A53 from TM1a. The TM1a swing introduces Y54 as a long-range seventh coordinating ligand. The seven-coordination of Mn^2+^ in the inward-open structure is less favored and more distorted, which may facilitate Mn^2+^ release. In contrast, the more favorable six- coordination in the outward-open and occluded structures could help energize the global conformational changes. The respective Mn^2+^-binding affinities reflect coordination chemistry: tighter Mn^2+^ binding at the orthosteric site correlates with more optimal coordination. Our metal- free structures indicate that after Mn^2+^ release to the cytosol, the protein resets to the outward- open state through the same occluded state. The residues in the gating networks are highly conserved, suggesting that their role is conserved across the Nramp family.

While the Mn^2+^ transport mechanism and associated conformational changes in DraNramp differ significantly from other LeuT-fold transporters^2, 17^, the presence of key residues defining the extracellular and intracellular gates is a common theme^28, 29, 44–49^. Comparing the gating networks described for these transporters with DraNramp, the positions and nature of the gating networks are generally not conserved, but two common themes emerge. First, opening (or closing) a particular vestibule often involves a pair of changes, as we observe for the intracellular vestibule in DraNramp (Fig. 6a). Such pairs of changes have been associated with “thin” and “thick” gates in Mhp1^50^ and LeuT^28^, that is, gates based on sidechain motions and helix motions, respectively. Second, some gating residue positions are shared, but the networks are not, indicating that different families have evolved analogous networks to stabilize equivalent conformations^28, 49^.

The accumulated structures of DraNramp also provide clues as to the thermodynamic landscape of the transport process (Fig. 6b). Interestingly, both metal-free and Mn^2+^-bound WT DraNramp crystallized in the occluded conformation, whereas the open conformations were achieved through conformation-locking^17^ or mutations of functionally important residues^18^. This occluded state more closely resembles an inward-open conformation^17^ and is an important intermediate in the Mn^2+^ transport cycle, with both local changes in coordination geometry and global changes in protein conformation in comparison to the outward-open state. Physiologically, we naïvely expect outward-open, rather than occluded, to be the preferred substrate-free conformation. However, the occluded or inward-open states of other LeuT-fold importers are often more stable, with changes in environmental conditions or the presence of substrate stabilizing specific states and lowering barriers to conformational transitions^51^. For example, SGLT1 and DraNramp require a negative membrane potential to transport substrates and this negative membrane potential stabilizes the outward-open state of SGLT1^21, 52^. Overall, our DraNramp structures suggest that the occluded state is most stable (at least in the absence of a membrane potential). Furthermore, our previous cysteine accessibility data indicate that the energy barriers between states are low enough for the protein to readily sample the outward- and inward-open states in cell membranes in the absence or presence of metal substrate^17, 18^.

Previous metal selectivity studies indicate that Nramps import different substrates with distinct mechanisms^2, 11, 15, 19, 21^. These differences correlate with chemical properties^53^ and preferred coordination chemistry^54, 55^ of these transition metals as observed in other transition metal binding proteins like the Psa permease^25, 38^, and cation diffusion facilitators (CDFs)^27^. That WT•Cd^2+^ retains the M230 sulfur as a coordinating ligand but excludes the D56 carboxylate can be rationalized by the fact that Cd^2+^ is a softer metal than Mn^2+^. This difference in coordination likely alters the pKa of nearby residues and water molecules to perturb the proton pathway such that DraNramp co-transports protons with Mn^2+^ but not Cd^2+^ (refs ^17, 21^), although our structures do not yet fully elucidate this mechanistic difference. Moreover, owing to its larger radius, Cd^2+^ tends to form weaker complexes than Mn^2+^ with comparable coordination numbers^54, 56^. The architecture of the orthosteric site in DraNramp appears to facilitate six- or seven-coordinated metal complexes, and thus most Cd^2+^-bound DraNramp conformations will be less stable than the Mn^2+^-bound states, as suggested by the ∼2-fold lower affinity for Cd^2+^ over Mn^2+^ at the orthosteric site. Our ITC data also indicate that binding of the toxic Cd^2+^ to DraNramp is less robust to perturbations compared to its physiological substrate Mn^2+^ and that Cd^2+^ only binds well to the inward-open state. The crystallographic data similarly suggests that inward-open is the most stable Cd^2+^-bound state (Fig. 6b), based on the following observations: (i) soaking crystals of occluded WT yielded inward-occluded WT•Cd^2+^, (ii) co-crystallization efforts yielded no other Cd^2+^-bound structures, and (iii) soaking Cd^2+^ into crystals of outward-locked G223W yielded very poor diffraction and no structures.

Overall, our data show that the orthosteric metal-binding site of DraNramp, conserved across all Nramps, is best suited to the physiological substrate Mn^2+^ (and likely the similar ion Fe^2+^). The distinct interactions of Nramps with Mn^2+^ and Cd^2+^ could be leveraged for the design of therapies for metal toxicity and prevention strategies for toxic metal accumulation in crops. These results also lay a foundation for future studies of how metal ion transporters like Nramps evolve their substrate selectivity, for example in response to different environmental conditions. Finally, the first complete set of structures with the same homolog, both in substrate-free and substrate-bound states, suggest a substrate-specific thermodynamic landscape of the transport cycle and provide a framework for future experiments and simulations to fully define this landscape, and for comparisons to other LeuT-fold transporters.

## Acknowledgements

We thank Arghya Deb for discussions of metal coordination chemistry, and previous and current members of the Gaudet lab for discussions and assistance, particularly Gerardo Zavala and Edward Lee for contributions to preliminary molecular dynamics analyses, and José Velilla for help with crystal soaking and fishing prior to data collection. This work was funded by NIGMS grant R01GM120996 (R.G.), a National Science Foundation (NSF) CAREER award MCB- 1942763 (A.S.), AstraZenaca and ASU-Mayo Foundation (E.W.), and the NSF-Simons Center for Mathematical and Statistical Analysis of Biology at Harvard (award number 1764269) and the Harvard Quantitative Biology Initiative (S.B.). Diffraction data reported in this study were collected at NE-CAT beamline 24IDC and GM/CA beamline 23IDB in the Advanced Photon Source. NE-CAT is funded by NIGMS grant P30 GM124165 and GM/CA is funded by the National Cancer Institute (ACB-12002) and the National Institute of General Medical Sciences (AGM-12006, P30GM138396). The Eiger 16M detector at GM/CA-XSD is funded by NIH grant S10 OD012289. The Advanced Photon Source is a U.S. Department of Energy Facility operated by Argonne National Laboratory under Contract No. DE-AC02-06CH11357. The molecular simulations used the Extreme Science and Engineering Discovery Environment (XSEDE) supported by NSF (ACI-1548562), and Oak Ridge Leadership Computing Facility, supported by the Office of Science, Department of Energy (DE-AC05-00OR22725).

## Author Contributions

RG and SR conceptualized the study. SR performed the crystallography, functional assays, and ITC experiments. EW and MS performed the MD simulations under the supervision of AS, SB analyzed the trajectories, and CZ and SB performed the sequence analyses. SR and RG wrote the manuscript, and all authors edited the manuscript.

## Online Methods

### Cloning and protein expression vectors

The DraNramp WT and mutant constructs were cloned into pET21-N8H^19^. All constructs used for crystallization had a truncation of 31 residues at the N-terminus (ΔN31, which does not impair metal transport(ref)), except for A47W which was full-length and transport deficient^18^. For proteoliposome-based transport assays, the full-length versions of each construct were used. For ITC, full-length DraNramp constructs were cloned into pET21-NStrep^19^ to avoid background signal from metals (Mn^2+^, Cd^2+^) binding to the His-tag. All mutations were introduced by site-directed mutagenesis using the Quikchange mutagenesis protocol (Stratagene) and confirmed by Sanger DNA sequencing.

### Protein expression

Protein expression was performed as previously described^19^. Briefly, transformed *Escherichia coli* C41(DE3) (Lucigen) cells were induced with 0.1 mM isopropyl-β-D- thiogalactopyranoside and cultured at 18°C for 16 h. Cell pellets from 10 L of culture were harvested and flash frozen in liquid nitrogen.

### Protein purification for crystallography

Cells were thawed and resuspended in 50 mL load buffer (20 mM sodium phosphate, pH 7.5, 55 mM imidazole pH 7.5, 500 mM NaCl, 10% (v/v) glycerol) supplemented with 1 mM PMSF, 1 mM benzamidine, 0.3 mg/mL DNAse I and 0.3 mg/mL lysozyme and lysed by sonication on ice (six cycles of 45 s with a Branson Sonifier 450 under duty cycle of 65% and output 10). Lysates were cleared by centrifuging for 20 min at 20,000 rpm (Beckman JA-20) and membranes pelleted from the supernatant by ultracentrifugation at 45,000 rpm (Beckman type 45Ti) for 70 min. Membranes were homogenized in 70 mL load buffer using a glass Potter-Elvehjem grinder, solubilized for 1 h in 1% (w/v) n-dodecyl-β-D-maltopyranoside (DDM), then ultracentrifuged at 35,000 (Beckman type 45Ti) for 35 min to remove insoluble debris. Pre-equilibrated Ni-Sepharose beads (2 mL; GE Healthcare) were incubated with the supernatant for 90 min at 4°C, then washed with 20 column volumes (CV) of each of the following buffers sequentially (i) load buffer containing 0.03 % DDM, (ii) load buffer containing 0.5% lauryl maltose neopentyl glycol (LMNG), and (iii) load buffer containing 0.1% LMNG. Protein was eluted in 20 mM sodium phosphate, pH 7.5, 450 mM imidazole pH 7.5, 500 mM NaCl, 10% (v/v) glycerol, 0.01% LMNG, concentrated to < 0.5 mL in a 50 kDa molecular weight cutoff (MWCO) centrifugal concentrator (EMD Millipore), and purified by size exclusion chromatography (SEC) using a Superdex S200 10/300 (GE Healthcare) pre-equilibrated with SEC buffer (10 mM HEPES pH 7.5, 150 mM NaCl, 0.003% LMNG). Peak protein fractions enriched in DraNramp were combined, concentrated to ∼25-40 mg/mL using a 50 kDa MWCO centrifugal concentrator, aliquoted and flash frozen in liquid nitrogen and stored at -80°C.

### Purification of DraNramp for ITC

To purify protein for ITC, harvested cells expressing strep-tagged DraNramp from 10 L of culture were resuspended in 50 mL of buffer W (100 mM Tris, pH 8.0, 150 mM NaCl), and membranes were isolated, homogenized and solubilized in 1% DDM as above. The supernatant was incubated with Strep-Tactin Superflow resin (3 mL; IBA) pre-equilibrated with buffer W + 0.03% DDM and washed with the following buffers sequentially: (i) 1 CV buffer W + 0.03% DDM, (ii) 2 CV buffer W + 0.5% LMNG and (iii) 2 CV buffer W + 0.1% LMNG. Protein was eluted with 3 CV buffer W + 0.01% LMNG + 2.5 mM desthiobiotin. The eluted protein was concentrated up to 2.5 mL using a 50 kDa MWCO centrifugal concentrator and buffer-exchanged into 150 mM NaCl, 10 mM HEPES, pH 7.5, and 0.003% LMNG using disposable PD-10 desalting columns (GE healthcare). Protein was concentrated to ∼ 2.5 mg/mL a 50 kDa MWCO centrifugal concentrator and flash frozen in liquid nitrogen and stored at -80°C.

### DraNramp crystallization

Crystallization of all constructs was performed using lipidic cubic phase (LCP). Protein was mixed with monoolein in 1:1.5 volume ratio using the syringe reconstitution method^17^. The protein bolus (60 nL) and 720 nL precipitant were dispensed onto custom-made 96 well glass sandwich plates using an NT8 drop-setting robot (Formulatrix). Metal-free crystals (WT) and metal supplemented (5 mM MnCl_2_) co-crystals (A47W•Mn^2+^, M230A•Mn^2+^, D296A•Mn^2+^) were grown in different precipitant conditions (Supplementary Table 1), harvested within 7-10 days (after reaching their optimal size of 30-40 μm rods) using mesh loops (MiTeGen) and flash-frozen in liquid nitrogen prior to data collection. Some structures (WT•Cd^2+^, WT•Mn^2+^, WT_soak_) were obtained by soaking WT DraNramp crystals grown in metal-free precipitant (Supplementary Table 1) for 7-10 days: The glass covering the wells was broken without disturbing the bolus and 2 μl soak solution (Supplementary Table 1) were added before resealing with a fresh siliconized glass coverslip, incubating overnight (16-18 h), then harvesting and flash freezing for data collection.

### X-ray diffraction data collection and processing

Diffraction data for structure determination and refinement were collected at beamlines 24-ID-C or 23-ID-B of the Advanced Photon Source at wavelengths of 0.984 Å or 1.033 Å, respectively. We used anomalous signals confirm the presence of metals (Mn^2+^ or Cd^2+^) in the binding site. For D296A•Mn^2+^ and A47W•Mn^2+^, the same data we used for structure refinement and collected at 1.033 Å, 0.984 Å, respectively, provided strong anomalous signal in the metal binding sites. For WT•Cd^2+^, we were able to collect data at 1.904 Å (near the low-energy boundary for the beamline), to maximize the anomalous signal. Locations of the crystals in the mesh loops were identified by grid scanning with a 20-μm beam at 10% transmission followed by data collection with a 10-μm beam at 15% transmission. Data were indexed in XDS^57^ and scaled in CCP4 AIMLESS (Version 7.0)^58, 59^. For datasets collected from several crystals, data from each crystal were independently indexed and integrated, then combined during scaling using CCP4 AIMLESS^58, 59^ to obtain complete datasets. Initial phases for all structures were determined by molecular replacement in PHENIX (Version 1.17.1-3660)^60^ using an occluded structure of DraNramp (PDB ID 6C3I chain A)^17^ as search model. Data statistics are listed in Table 1 and Supplementary Tables 2 and 4.

### Model building, refinement, and analysis

Models were built in COOT (Version 7.0)^61^ and refined in PHENIX^60^, with macrocycles including reciprocal space, TLS groups, and individual B-factor refinement, and optimization of the X-ray/stereochemistry and X-ray/ADP weights. For WT•Cd^2+^, ‘anomalous group refinement’ was used to improve the fit to density of the Cd^2+^ ions, with Cd^2+^ as an ‘anomalous group’ with the reference f’ and f” values suggested by phenix.form.factor (-0.462 and 2.132 respectively). Ligand restraints for monoolein and spermidine were generated in Phenix.elbow with automatic geometry optimization. All structures contain one protein molecule in the asymmetric unit. The final structures span from residues 45–48 to residues 433–436, except that residues 240–249 and 240–247 were not modeled in D296•Mn^2+^ and WT•Cd^2+^, respectively, because of lack of interpretable electron density map. Model refinement statistics are listed in Table 1 and Supplementary Tables 2 and 4. Pairwise RMSD for all structures are listed in Supplementary Table 3. Anomalous difference Fourier maps for Mn^2+^ (D296A•Mn^2+^ and A47W•Mn^2+^) and Cd^2+^ (WT•Cd^2+^) were generated in phenix.maps using a high-resolution cutoff of 3.5–4.5 Å. Polder maps omitting the metal ions were generated in phenix.polder to appropriately define the coordination sphere for Mn^2+^ and Cd^2+^ in all structures. All software were provided by SBGrid^62^.

### Metal binding measurements using ITC

ITC experiments were performed using MicroCal iTC200 (GE Healthcare) to determine the affinity and thermodynamic parameters of binding of divalent metals (Mn^2+^, Cd^2+^ and Mg^2+^) to DraNramp (WT and its mutants)^63, 64^. All protein and metal solutions were prepared in ITC buffer (150 mM NaCl, 10 mM HEPES, pH 7.5, and 0.003% LMNG). The sample cell containing 25 μM protein was titrated with 20 2-μL injections of 6 mM metal, with an interval of 120 sec between each successive 5-s injection, with a 750-rpm stirring rate at 25°C. To nullify the heat of dilution, the data from titration of a metal solution into ITC buffer (‘buffer blank’ runs) were subtracted from the metal-protein titration curves prior to model fitting. Data were fitted and analyzed using a one- or two-site model (depending on the number of metal binding sites for each construct) with Origin 7 software. The mean ± SEM from 2-3 repeats for each sample are reported in Supplementary Table 7. Detailed thermodynamic parameters for each run are reported in Supplementary Tables 8 and 9.

### Proteoliposome-based *in vitro* transport assays

Protein purification, liposome preparation, and metal transport assays were performed as described^19, 21^.

### Sequence alignments

We used 92 Nramp sequences from Pfam^65^ to build a seed alignment using MUSCLE^66^. We collected 15,451 sequences from Uniprot^67^ using HMMER^68^. We used HMMER’s hmmalign, with a hidden Markov model profile from the seed alignment as an input, to align all 15,451 sequences. We applied filters to retain sequences 400-600 residues in length and sequences with under 90% pairwise sequence identity, respectively. The final alignment contains 6712 sequences and is well aligned at biologically relevant residues. A maximum-likelihood phylogenetic tree was generated via RAxML-NG^69^ using the LG substitution model^70^, with the likeliest final tree selected from ten parallel optimization trials. The canonical Nramp clade in this tree was identified based on conservation of the "DPGN" and "MPH" motifs in transmembrane helices 1 and 6, respectively, and contained 3796 sequences. The Nramp-related magnesium transporters were used to root the canonical Nramp phylogeny. Sequence analysis was done with Biopython^71^ and sequence logos were generated with logomaker^72^ using a "chemistry" color scheme inspired by Weblogo^73^.

### Molecular Dynamics simulation

Molecular dynamics (MD) simulations were initialized from three high-resolution structures of DraNramp: the outward-open G223W•Mn^2+^ structure (6BU5) with Mn^2+^ removed and W223 mutated back to the native glycine residue *in silico*, the inward-open WT•Cd^2+^ structure with Cd^2+^ removed, and the inward-occluded WT structure. Crystallographic waters were retained, and protonation states of key titratable residues were selected with PROPKA^74, 75^ assuming a pH of 5.0 for residues exposed to external solvent and a pH of 7.0 for residues exposed to cytosol, a condition under which DraNramp exhibits high activity. All structures were oriented in the membrane with the PPM web server and membrane systems were prepared with CHARMM- GUI^76, 77^. A POPC membrane of surface area 99 x 99 Å was constructed in the XY plane around the protein^78^, the system was solvated in a 100 x 100 x 100 Å^3^ rectangular box using TIP3 waters and electronically neutralized using potassium and chlorine ions at an overall concentration of 150 mM. The overall system size was approximately 103,000 atoms.

All-atom simulations were run using GPU-accelerated NAMD^79^ and the CHARMM36m forcefield^80^. Prior to simulation, the energy of each system was minimized for 10,000 steps using a conjugate gradient and line search algorithm native to NAMD. To improve simulation stability, the system was initially equilibrated using an NVT-ensemble with harmonic restraints placed on protein and lipid heavy atoms. The harmonic restraints were then incrementally relaxed over a period of 675 ps according to established CHARMM-GUI protocols^77^. The system was then simulated at a constant pressure, utilizing the Langevin piston method to maintain 1 atm at 303.15 K, from anywhere between 617 to 1176 ns depending on the starting conformation.

Simulations were performed using periodic boundary conditions and a time step of 3.0 fs with all bonds to hydrogens being constrained. Large integration timesteps were enabled by employing hydrogen mass repartitioning^81^. Long-range electrostatic interactions were calculated using the particle mesh Ewald (PME) method with nonbonded interactions being cut off at 12 Å. Each simulation was performed in duplicate resulting in approximately 2 µs of total sampling for each system. Simulations are summarized in Supplementary Table 10.

RMSD, residue distance, and dihedral analyses were performed using mdtraj version 1.9.8^82^. For distance analysis, the minimum interatomic distances were identified between the specified residues across each frame. For water analysis, simulations were centered and wrapped in VMD and a Tcl script was used to produce water density maps. Contour maps of these densities were then visualized in PyMOL. Water occupancies of sites coordinated by specific residues were also calculated in an alignment-agnostic manner by determining for each frame in each simulation whether a water was present within 2.5 Å of both specified residues. Distinct rotamers were identified from dihedrals using spectral clustering as implemented in scikit-learn version 1.0.

### Data Availability

Atomic coordinates and structure factors for the crystal structures reported in this work have been deposited to the Protein Data Bank under accession numbers 8E5S (WT), 8E5V (WT_soak_), 8E60 (WT•Mn^2+^), 8E6H (A47W•Mn^2+^), 8E6I (M230A•Mn^2+^), 8E6L (D296A•Mn^2+^), 8E6M (WT•Cd^2+^), and 8E6N (re-refined G223W•Mn^2+^). Corresponding X-ray diffraction images have been deposited to the SBGrid Data Bank under the respective accession numbers 962 (doi:10.15785/SBGRID/962), 963 (doi:10.15785/SBGRID/963), 964 (doi:10.15785/SBGRID/964), 966 (doi:10.15785/SBGRID/966), 967 (doi:10.15785/SBGRID/967), 968 (doi:10.15785/SBGRID/968), 969 (doi:10.15785/SBGRID/969), and previously deposited 564 (doi:10.15785/ SBGRID/564).

---

**Extended Data Figure 1.**
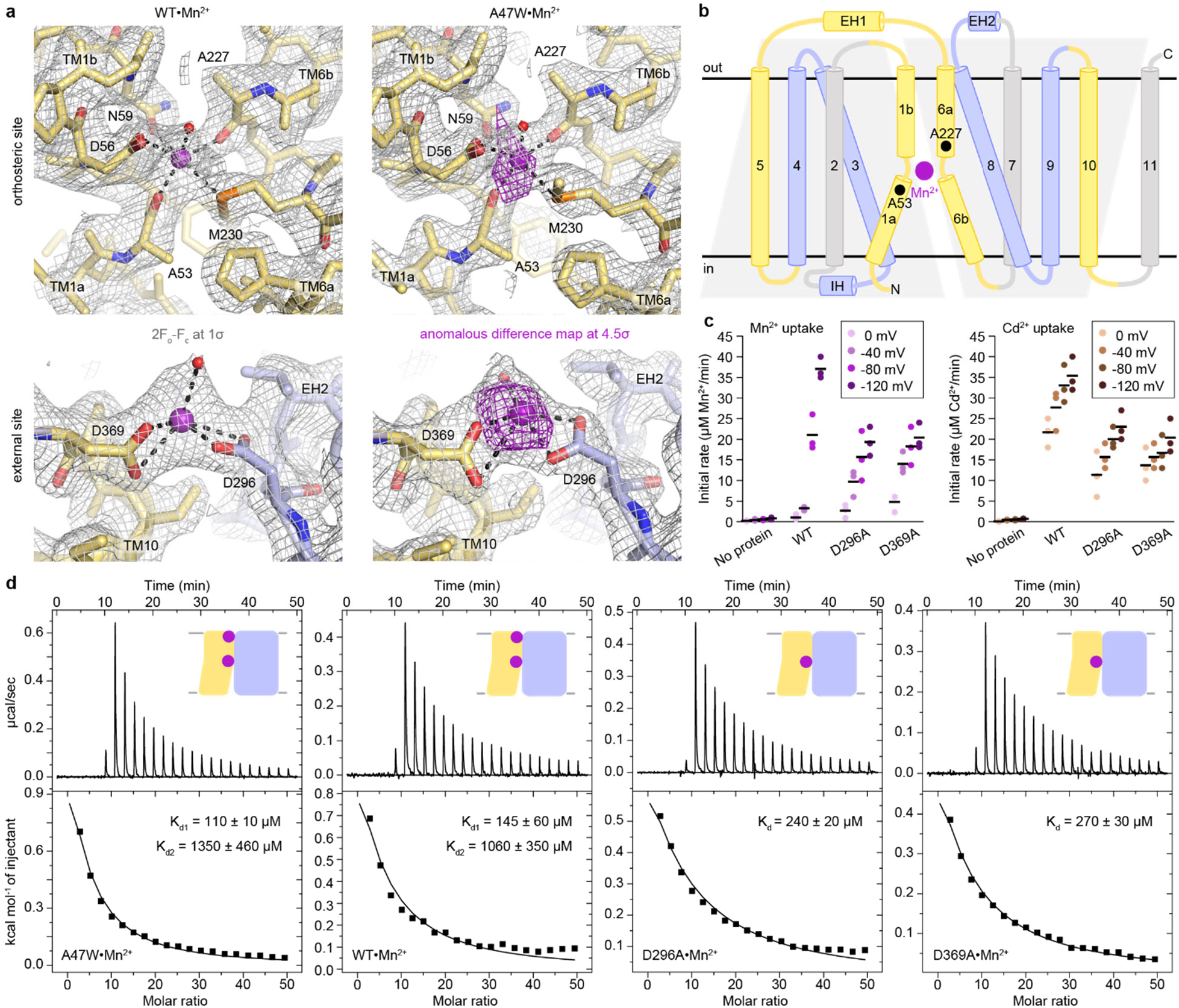
Structure and affinity of Mn^2+^ binding at the orthosteric and external sites of DraNramp. (a) 2F_o_-F_c_ (grey mesh; 1σ) maps and peaks from anomalous difference Fourier (magenta mesh; 4.5σ) calculated from the WT•Mn^2+^ and A47W•Mn^2+^ structures, respectively show Mn^2+^ bound at two sites, the orthosteric site (top) and an external site coordinated by D296 and D369 near the N-termini of EH2 and TM10, respectively (bottom). (b) Topology diagram showing the secondary structure organization of DraNramp with its characteristic LeuT fold, where TMs 1–5 and 6–10 form two pseudosymmetric inverted repeats. One intracellular helix, IH, and two extracellular helices, EH1 and EH2, connect TMs 2-3, 5-6, and 7-8, respectively. Black spheres indicate A53 in TM1a and A227 in TM6a, the two Mn^2+^-coordinating backbone carbonyls in the occluded state of DraNramp, one in each inverted repeat. Mn^2+^ is shown as magenta sphere. (c) Initial metal uptake rates for DraNramp mutants at membrane potentials ranging from ΔΨ = 0 to −120 mV (n = 3; black bars are the mean values). The metal ion concentration was 750 μM, and the pH was 7 on both sides of the membrane. D296A and D369A moderately reduced the initial transport rate at high membrane potentials. The overall trends are similar for both metals. Mn^2+^ transport showed higher voltage dependence than Cd^2+^ transport. Corresponding time traces are plotted in Supplementary Figure 1. (d) ITC measurements of Mn^2+^ binding to A47W (which behaves like WT), WT (reproduced here from Figure 1 for comparison), and D296A and D369A. Either of the external site mutations are best fit as Mn^2+^ binding to a single site, and the resulting K_d_ values are most similar to K_d1_ of WT, indicating that the orthosteric site has higher Mn^2+^ binding affinity compared to the external site.

**Extended Data Figure 2.**
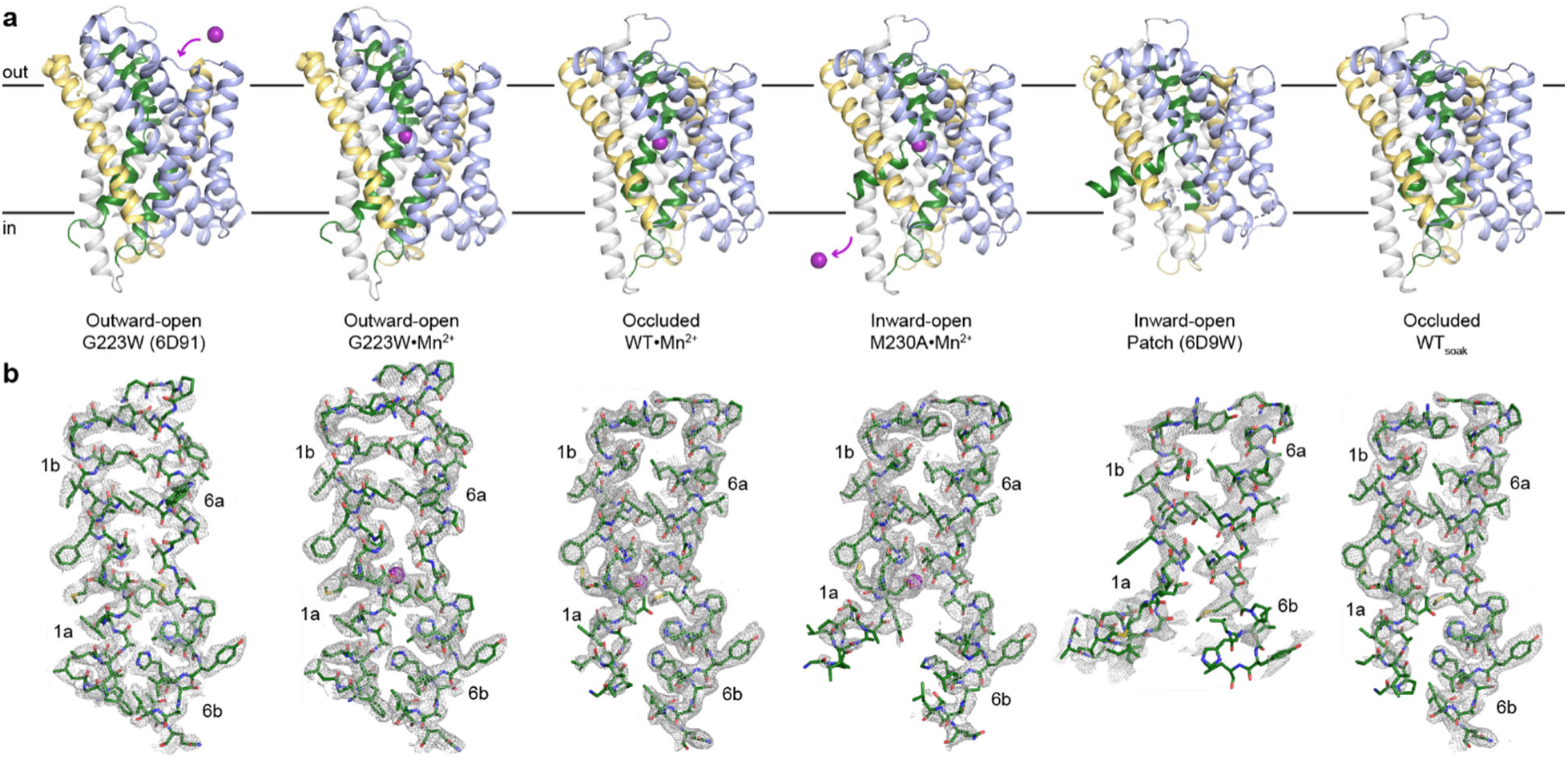
Structures of different conformational states of the DraNramp Mn^2+^ transport cycle. (a) Crystal structures of each state of DraNramp. TMs 1 and 6 are colored green, TMs 5 and 10 are colored pale yellow, TMs 2, 7 and 11 in gray and TMs 3, 4, 8, and 9 are light blue. The Mn^2+^ ion in the orthosteric site, when present, is shown as a magenta sphere. (b) Corresponding 2F_o_-F_c_ maps contoured at 1σ for TMs 1 and 6 of each structure illustrated in **a**.

**Extended Data Figure 3.**
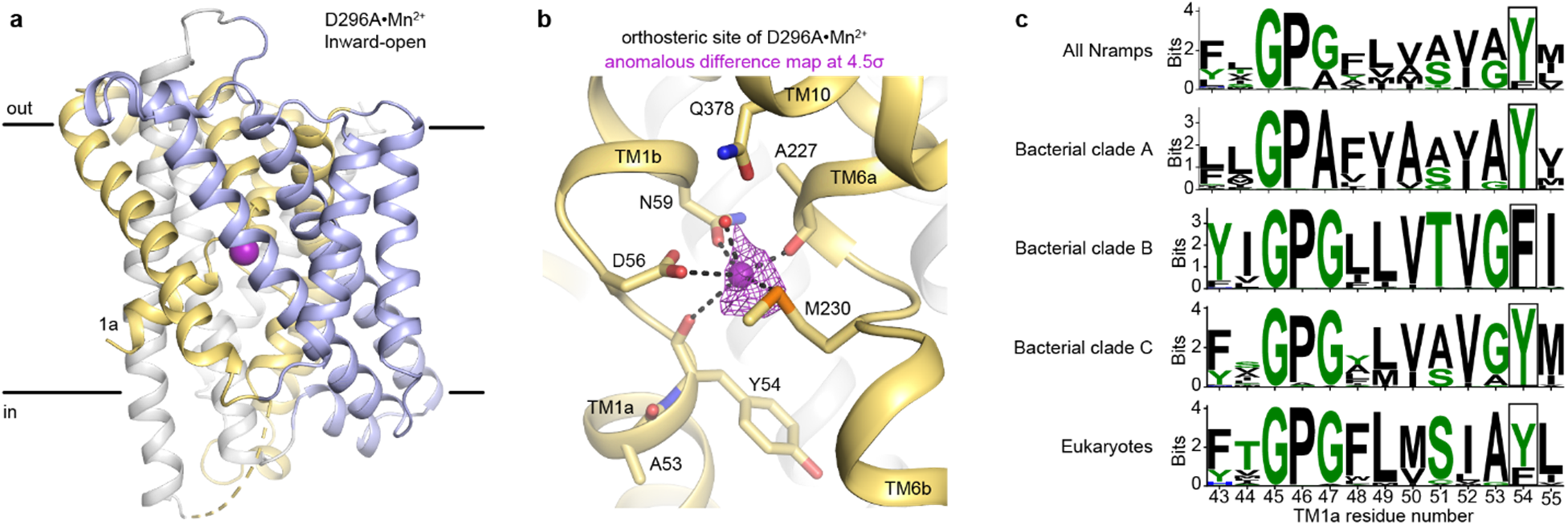
Binding of Mn^2+^ at the orthosteric site and conservation of an aromatic residue at position 54 involved in opening of the inner gate. (a) Cartoon representation of the inward-open structure of D296A•Mn^2+^, which is nearly identical to the M230A•Mn^2+^ inward-open structure (Cα RMSD of 0.38 Å). (b) Coordination sphere of the orthosteric Mn^2+^ ion in the D296A structure is nearly the same as in M230A Mn^2+^-bound structure except for the sulfur of M230 replacing a water seen in M230A. We do not observe a bound water to complete the coordination sphere of D296A, likely because the resolution of the structure is lower (2.52 Å for M230A•Mn^2+^ vs. 3.12 Å for D296A•Mn^2+^), otherwise the structures are analogous. The peak from the anomalous difference Fourier map (magenta mesh; 4.5σ) calculated from a D296A•Mn^2+^ crystal confirms a bound Mn^2+^ in this inward-open state. (c) Sequence logos highlighting that Y54 in TM1a is 80% conserved in all Nramps (3762 sequences), 100% conserved in bacterial clades A and C, but replaced by a phenylalanine in clade B. Eukaryotic Nramps have either tyrosine or phenylalanine at the corresponding position. Residue coloring is based on ‘chemistry’ coloring scheme of Weblogo^73^. Clades are defined as in Supplementary Fig. 2b.

**Extended Data Figure 4.**
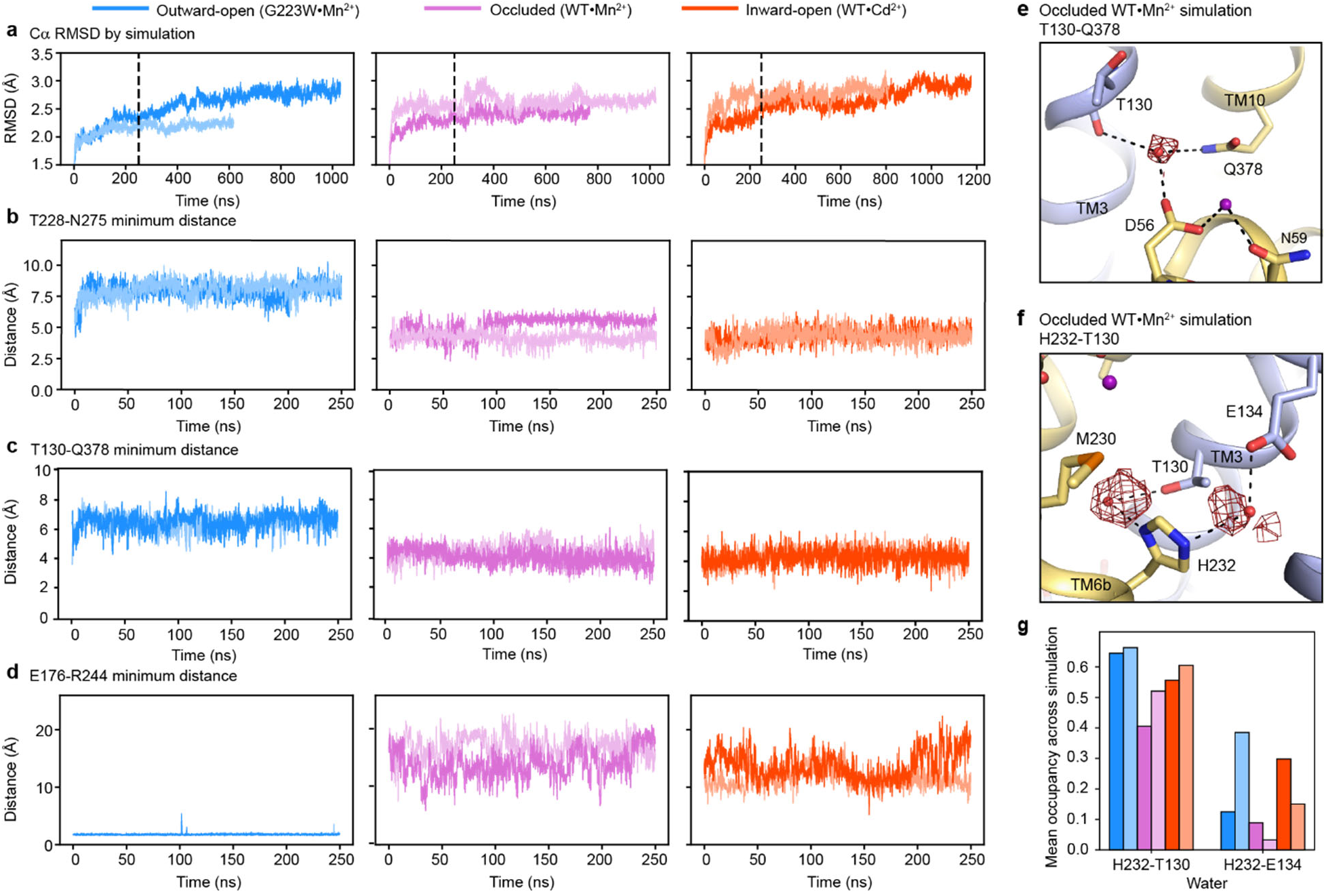
Structural stability of gating residue networks of the Mn^2+^ transport cycle. (a-d) Time-course analyses of MD simulations starting in each of the three major conformations, showing replicates in different shades. (a) RMSD, calculated over all Cα atoms relative to the starting structure. Because some simulations experience significant conformational changes after the first 250 ns (dashed line), we focused on this initial range for subsequent analyses. (b-c) Plots of the minimum distance between T228 and N275 (b) and T130 and Q378 (c) in the outer vestibule show that these interactions are stable in the inward-occluded and inward-open simulations but do not form in the outward-open simulations. (d) Likewise, simulations show that E176 and R244, located in the inner vestibule on TM5 and TM7, respectively, stably interact in the outward-open conformation, but not the occluded or inward-open conformations. (e-f) Representative 50% contour maps of water density calculated from a simulation starting the occluded conformation. (e) The water bridging T130, Q378 and D56 persists throughout most the simulation, as do (f) the waters coordinated by H232 and T130 and H232 and E134, suggesting that all these waters-mediated interactions are robust. (g) A water is coordinated by H232 and T130 in 40-60% of frames across all simulations. A water is also coordinated by H232 and E134 but less frequently, in 2-40% of frames.

**Extended Data Figure 5.**
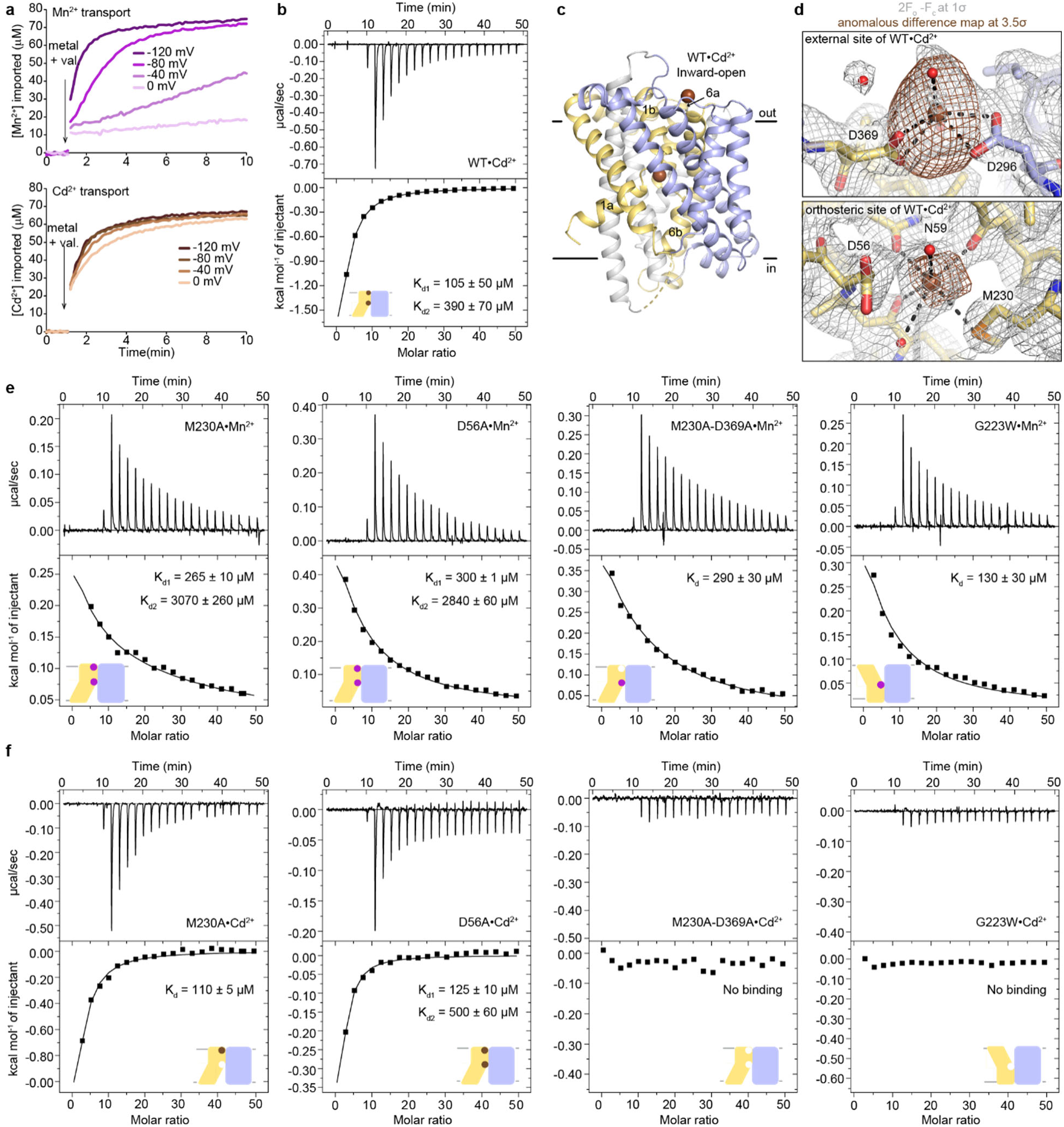
Analysis of Cd^2+^ and Mn^2+^ binding to DraNramp reveal differences. (a) Proteoliposome-based assay with WT DraNramp showed higher voltage dependence for Mn^2+^ transport compared to Cd^2+^, with substantial transport of only Cd^2+^ at 0 mV. (b) ITC of WT DraNramp binding to Cd^2+^. The data show an exothermic mode of binding and are fit using a two-binding-site model (K_d1_ = 105 ± 50 µM, K_d2_ = 390 ± 70 µM). By comparing ITC results from binding of Cd^2+^ to WT and DraNramp constructs with mutations at the external site (panel e), we assigned a lower affinity (K_d2_) to the orthosteric site. (c) Cartoon representation of WT•Cd^2+^, with Cd^2+^ ions as brown spheres. The positions of TM1 and TM6, especially the upward swing of TM1a, confirms the inward-open conformation. (d) Peaks from the anomalous difference Fourier map (brown mesh; 3.5σ) and electron density from the 2F_o_-F_c_ map (grey mesh; 1σ) from WT•Cd^2+^ show that Cd^2+^ binds at both the external and orthosteric sites. (e-f) ITC measurements comparing the affinity of various DraNramp constructs for Mn^2+^ (e) or Cd^2+^ (f). As described in the legend to Figure 5d, the M230A mutation at the orthosteric site impacts Cd^2+^ binding more than Mn^2+^. In the G223W outward-locked construct, the external-site ligands are far apart because of the opening of the extracellular vestibule (Supplementary Figure 2a) but the orthosteric site is intact. Mn^2+^ binds at the orthosteric site in G223W, but Cd^2+^ binding is impaired compared to WT. Mn^2+^- and Cd^2+^-binding isotherms are endothermic and exothermic, respectively.

**Extended Data Figure 6.**
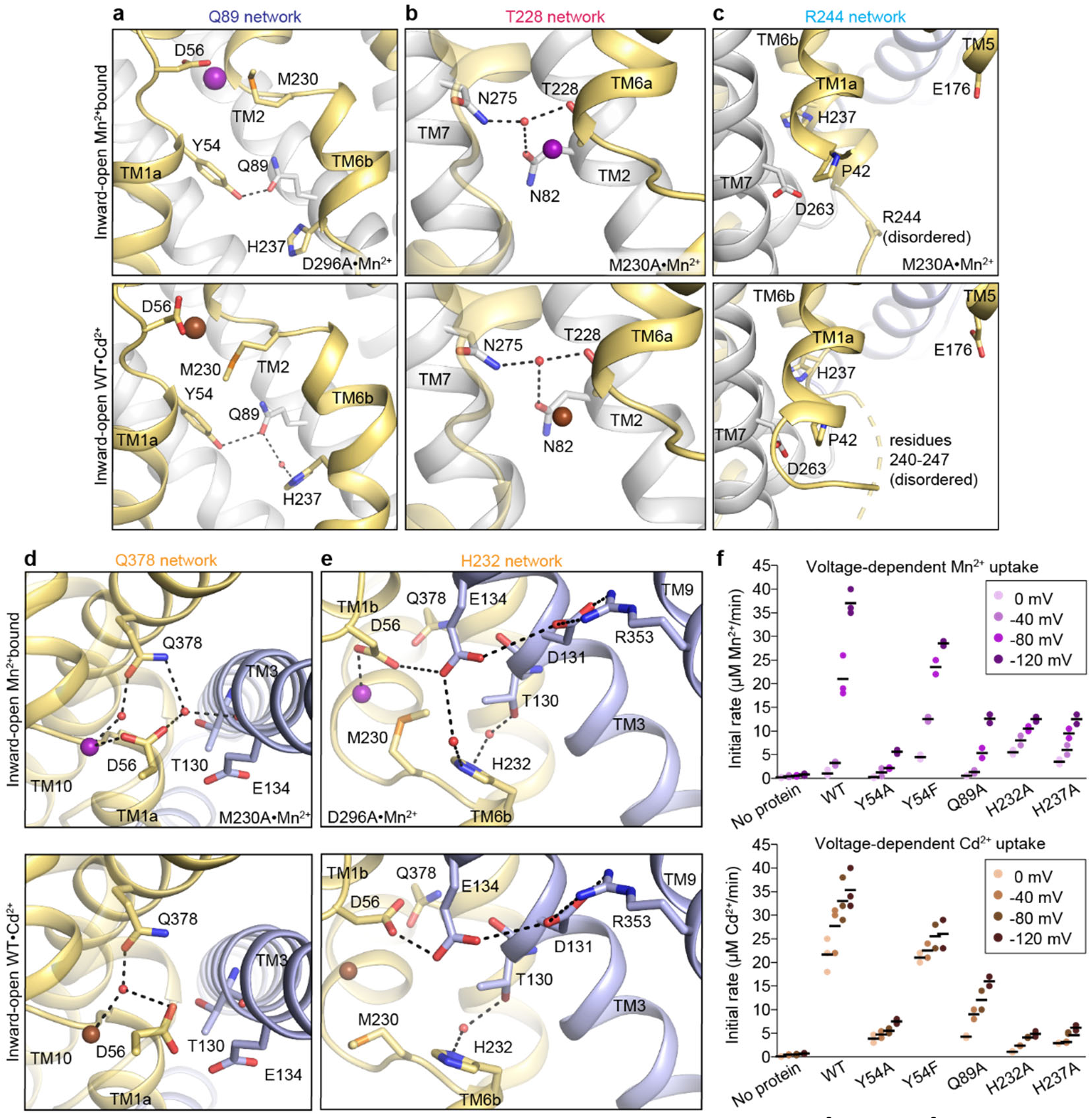
Comparison of the gating networks in the inward-open Mn^2+^ and Cd^2+^-bound structures. (a-e) Structural comparisons of the five networks of polar residues with Mn^2+^ (top; M230A•Mn^2+^ or D296A•Mn^2+^ as indicated) or Cd^2+^ (WT•Cd^2+^; bottom) bound at the orthosteric site. Inward-open conformations of Mn^2+^ and Cd^2+^ structures show identical arrangement of residues for the Q89 (a), T228 (b), and R244 (c) networks. In the outer-gate Q378 network (d), D56 does not coordinate the Cd^2+^ directly, but instead coordinates a water that interacts with Cd^2+^ and Q378. Due to its reorientation, D56 does not hydrogen-bond to T130. The H232 network (e) is conserved except that the interaction between E134 and D56 takes a different orientation in the Cd^2+^-bound structure. Some of the top (Mn^2+^-bound) panels are reproduced from Figure 4 for ease of comparison. (f) Initial metal uptake rates for DraNramp variants at different membrane potential (ΔΨ = 0 to −120 mV; n = 2-3; black bars are the mean values). The metal ion concentration was 750 μM, and the pH 7 on both sides of the membrane. Y54A, H232A, H237A, and Q89A strongly reduced the initial transport rate, whereas Y54F only moderately reduced it. The overall trends are similar for both metals. Mn^2+^ transport showed higher voltage dependence compared to Cd^2+^. Mn^2+^ data from Figure 3 are replotted here for ease of comparison. Corresponding time traces are plotted in Supplementary Figure 1.

**Supplementary Table 1.** Construct, precipitant, and soaking solutions used for each structure

| Structure<br>(Construct•substrate)<sup>a</sup> | Precipitant solution | Soaking solution<sup>c</sup> |
| --- | --- | --- |
| ΔN31-WT (no metal) | 28% PEG 400<br>0.1 M MES (pH 6.7)<br>50 mM succinic acid (pH 6.0)<br>5 mM spermidine (pH 7.0) | |
| ΔN31-WT<sub>soak</sub> | 28% PEG 400<br>0.1 M MES (pH 6.3)<br>50 mM succinic acid (pH 6.0)<br>5 mM spermidine (pH 7.0) | 28% PEG 400<br>0.1 M MES (pH 6.5)<br>50 mM succinic acid (pH 6.0)<br>20 mM spermidine (pH 7.0) |
| ΔN31-WT•Mn<sup>2+</sup> | 24% PEG 400<br>0.1 M MES (pH 5.9)<br>50 mM succinic acid (pH 6.0)<br>20 mM spermidine (pH 7.0) | 28% PEG 400<br>0.1 M MES (pH 6.5)<br>50 mM succinic acid (pH 6.0)<br>20 mM spermidine (pH 7.0)<br>2 mM MnCl<sub>2</sub> |
| ΔN31-WT•Cd<sup>2+</sup> | 24% PEG 400<br>0.1 M MES (pH 6.1)<br>50 mM succinic acid (pH 6.0)<br>20 mM spermidine (pH 7.0) | 28% PEG 400<br>0.1 M MES (pH 6.5)<br>50 mM succinic acid (pH 6.0)<br>20 mM spermidine (pH 7.0)<br>2 mM CdCl<sub>2</sub> |
| A47W•Mn<sup>2+</sup> (b) | 20% PEG MME 550<br>0.1 M HEPES (pH 7.2)<br>0.25 M NaCl<br>10 mM MnCl<sub>2</sub> |  |
| ΔN31-M230A•Mn<sup>2+</sup> (b) | 32% PEG 400<br>0.1 M MES (pH 6.5)<br>50 mM succinic acid (pH 6.0)<br>20 mM spermidine (pH 7.0)<br>10 mM MnCl<sub>2</sub> | |
| ΔN31-D296A•Mn<sup>2+</sup> (b) | 28% PEG 400<br>0.1 M HEPES (pH 6.8)<br>0.1 M NaCl<br>10 mM MnCl<sub>2</sub> | |

<sup>a</sup> ΔN31 refers to a deletion of the N-terminal 31 residues. For simplicity, the “ ΔN31” annotation is listed here, but omitted elsewhere in the text.
<sup>b</sup> Proteins were premixed with 5 mM MnCl<sub>2</sub> prior to setting up crystallization trials in precipitants spiked with 10 mM MnCl<sub>2</sub>.
<sup>c</sup> Crystals of ‘ ΔN31-WT (no metal)’ were soaked in the listed soaking solution overnight prior to harvesting.

**Supplementary Table 2.** Data collection and refinement statistics for the four supporting new DraNramp structures

| Structure | WT | A47W•Mn<sup>2+</sup> | D296A•Mn<sup>2+</sup> | G223W•Mn<sup>2+</sup> (a) |
| --- | --- | --- | --- | --- |
| Conformation | Occluded | Occluded | Inward open | Outward open |
| Bound substrate | none | Mn<sup>2+</sup> | Mn<sup>2+</sup> | Mn<sup>2+</sup> |
| PDB ID | 8E5S | 8E6H | 8E6L | 8E6N |
| **Data Collection** |  |  |  |  |
| Beamline | NECAT 24IDC | NECAT 24IDC | GMCA 23IDB | NECAT 24IDC |
| Wavelength (Å) | 0.984 | 0.984 | 1.033 | 0.979 |
| Resolution range (Å) | 41.06-2.38 (2.46-2.38) | 45.35-2.39 (2.47-2.39) | 45.32-3.12 (3.23-3.12) | 39.19-2.40 (2.49-2.40) |
| Space group | P 2 21 21 | P 2 21 21 | P 2 21 21 | C1 21 |
| Unit cell (*a*, *b*, *c*) | 58.66, 70.85, 98.34 | 58.98, 70.93, 98.57 | 58.51, 71.64, 98.89 | 105.76, 80.39, 51.75 |
| Unit cell ($\alpha$, $\beta$, $\gamma$) | 90, 90, 90 | 90, 90, 90 | 90, 90, 90 | 90, 94.72, 90 |
| Number of crystals | 1 | 1 | 3 | ~15 |
| Total reflections | 74541 (7226) | 61843 (6302) | 27350 (2837) | 49321 |
| Unique reflections | 15762 (1607) | 16817 (1653) | 7004 (687) | 13962 (621) |
| Redundancy | 4.7 (4.5) | 3.7 (3.8) | 3.9 (4.1) | 3.5 (1.7) |
| Completeness (%) | 91.66 (93.38) | 98.57 (99.52) | 89.46 (90.75) | 82.1* (36.4) |
| Mean *I*/$\sigma$(*I*) | 9.83 (1.60) | 7.92 (1.03) | 6.71 (1.49) | 6.2 (1.6) |
| Wilson *B*-factor | 36.73 | 40.33 | 63.48 | 58.94 |
| *R*<sub>merge</sub> | 0.192 (1.057) | 0.163 (1.371) | 0.342 (1.216) | 0.186 |
| *R*<sub>meas</sub> | 0.218 (1.189) | 0.191 (1.588) | 0.386 (1.380) | 0.21 |
| *R*<sub>pin</sub> | 0.099 (0.528) | 0.097 (0.789) | 0.173 (0.634) | 0.095 |
| CC1/2 | 0.99 (0.51) | 0.98 (0.41) | 0.80 (0.53) | 0.98 (0.58) |
| **Refinement** |  |  |  |  |
| Resolution range (Å) | 41.06-2.38 (2.46-2.38) | 45.35-2.39 (2.47-2.39) | 45.32-3.12 (3.23-3.12) | 39.19-2.40 (2.49-2.40) |
| No. reflections | 15665 (1579) | 16749 (1653) | 6999 (687) | 13963 (621) |
| No. reflections in *R*<sub>free</sub> | 1565 (159) | 1673 (165) | 910 (90) | 701 (33) |
| *R*<sub>work</sub> | 0.206 (0.264) | 0.205 (0.299) | 0.219 (0.287) | 0.223 (0.276) |
| *R*<sub>free</sub> | 0.248 (0.312) | 0.246 (0.371) | 0.272 (0.352) | 0.271 (0.337) |
| Number of atoms | 3403 | 3483 | 3205 | 3310 |
| Protein | 2922 | 2939 | 2861 | 3012 |
| Ligand | 434 | 469 | 323 | 244 |
| Water | 47 | 75 | 21 | 54 |
| Protein Residues | 392 | 392 | 385 | 398 |
| Ramachandran plot |  |  |  |  |
| Favored (%) | 98.72 | 99.74 | 98.16 | 97.73 |
| Allowed (%) | 1.28 | 0.26 | 1.84 | 2.27 |
| Outliers (%) | 0 | 0 | 0 | 0 |
| Rotamer outliers (%) | 0.34 | 1.00 | 0.69 | 1.29 |
| Clashscore | 7.21 | 6.38 | 7.04 | 6.48 |
| RMS (bonds) | 0.003 | 0.002 | 0.002 | 0.002 |
| RMS (angles) | 0.49 | 0.43 | 0.45 | 0.44 |
| Average *B*-factor | 48.09 | 50.38 | 68.03 | 75.00 |
| Protein | 46.20 | 47.30 | 68.42 | 74.11 |
| Ligand | 61.54 | 70.13 | 65.22 | 88.03 |
| Water | 41.47 | 47.56 | 58.65 | 65.83 |
| No. of TLS groups | 7 | 3 | 8 | 5 |

Values in parentheses are for highest-resolution shell. Data for D296A•Mn<sup>2+</sup> merge reflections from multiple crystals. Data for the other structures were obtained from a single crystal.
<sup>a</sup> G223W•Mn<sup>2+</sup> has been re-refined from PDB ID: 6BU5, the previously published data collection statistics<sup>1</sup> are reproduced here for completeness.

**Supplementary Table 3.** C*α* RMSD in Å for all DraNramp structure pairs (number of aligned residues in parentheses)

| Conformation |  | Outward open |  | Occluded |  |  |  |  | Inward open |  |  |
| --- | --- | --- | --- | --- | --- | --- | --- | --- | --- | --- | --- |
|  | Structure | G223W<sup>(a)</sup> | G223W•Mn<sup>2+</sup> | WT<sub>soak</sub> | WT | WT•Mn<sup>2+</sup> | A47W•Mn<sup>2+</sup> | Patch<sup>(a,b)</sup> | M230A•Mn<sup>2+</sup> | D296A•Mn<sup>2+</sup><sup>(b)</sup> | WT•Cd<sup>2+</sup> |
| Outward open | G223W<sup>(a)</sup> |  | 0.97<br>(396) | 2.29<br>(359) | 2.29<br>(356) | 2.33<br>(361) | 2.39<br>(359) | 2.57<br>(320) | 2.45<br>(356) | 2.38<br>(344) | 2.32<br>(347) |
|  | G223W•Mn<sup>2+</sup> | 0.97<br>(396) |  | 2.34<br>(348) | 2.38<br>(350) | 2.39<br>(351) | 2.47<br>(353) | 2.67<br>(320) | 2.62<br>(355) | 2.47<br>(339) | 2.47<br>(398) |
| Occluded | WT<sub>soak</sub> | 2.29<br>(359) | 2.34<br>(348) |  | 0.38<br>(391) | 0.20<br>(392) | 0.43<br>(388) | 1.54<br>(328) | 0.56<br>(386) | 0.61<br>(366) | 0.77<br>(372) |
|  | WT | 2.29<br>(356) | 2.38<br>(350) | 0.38<br>(391) |  | 0.39<br>(392) | 0.42<br>(391) | 1.69<br>(329) | 0.71<br>(387) | 0.82<br>(367) | 0.95<br>(374) |
|  | WT•Mn<sup>2+</sup> | 2.33<br>(361) | 2.39<br>(351) | 0.20<br>(392) | 0.39<br>(392) |  | 0.47<br>(389) | 1.53<br>(328) | 0.59<br>(387) | 0.65<br>(367) | 0.79<br>(373) |
|  | A47W•Mn<sup>2+</sup> | 2.39<br>(359) | 2.47<br>(353) | 0.43<br>(388) | 0.42<br>(391) | 0.47<br>(389) |  | 1.59<br>(326) | 0.59<br>(383) | 0.61<br>(363) | 0.60<br>(368) |
| Inward open | Patch<sup>(a,b,c)</sup> | 2.57<br>(320) | 2.67<br>(320) | 1.54<br>(328) | 1.69<br>(329) | 1.53<br>(328) | 1.59<br>(326) |  | 1.59<br>(339) | 1.45<br>(340) | 1.52<br>(341) |
|  | M230A•Mn<sup>2+</sup> | 2.45<br>(356) | 2.62<br>(355) | 0.56<br>(386) | 0.71<br>(387) | 0.59<br>(387) | 0.59<br>(383) | 1.59<br>(339) |  | 0.38<br>(372) | 0.48<br>(376) |
|  | D296A•Mn<sup>2+</sup><sup>(b,c)</sup> | 2.38<br>(344) | 2.47<br>(339) | 0.61<br>(366) | 0.82<br>(367) | 0.65<br>(367) | 0.61<br>(363) | 1.45<br>(340) | 0.38<br>(372) |  | 0.47<br>(384) |
|  | WT•Cd<sup>2+</sup><sup>(c)</sup> | 2.32<br>(347) | 2.47<br>(398) | 0.77<br>(372) | 0.95<br>(374) | 0.79<br>(373) | 0.60<br>(368) | 1.52<br>(341) | 0.48<br>(376) | 0.47<br>(384) |  |

<sup>a</sup> Previously published structures (G223W is PDB ID: 6D91; Patch is PDB ID: 6D9W)
<sup>b</sup> Lower-resolution structures.
<sup>c</sup> Structures with unmodeled loops.

**Supplementary Table 4.** Data collection for anomalous maps

|  | A47W•Mn<sup>2+</sup><br>Occluded | D296A•Mn<sup>2+</sup><br>Inward open | WT•Cd<sup>2+</sup><br>Inward open |
| --- | --- | --- | --- |
| Beamline | NECAT 24IDC | GMCA 23IDB | NECAT 24IDC |
| Wavelength (Å) | 0.984 | 1.033 | 1.904 |
| Resolution range (Å) | 45.35-2.38 (2.46-2.38) | 45.32-3.12 (3.23- 3.12) | 40.28-2.82 (2.92-2.82) |
| Space group | P 2 21 21 | P 2 21 21 | P 2 21 21 |
| Unit cell (*a*, *b*, *c*) | 58.98, 70.93, 98.57 | 58.51, 71.64, 98.89 | 58.54, 70.72, 98.00 |
| Unit cell ($\alpha$, $\beta$, $\gamma$) | 90, 90, 90 | 90, 90, 90 | 90, 90, 90 |
| Total reflections | 62633 (6419) | 27350 (2837) | 32558 (3197) |
| Unique reflections | 17027 (1685) | 7000 (688) | 10017 (989) |
| Redundancy | 3.7 (3.8) | 3.9 (4.1) | 3.3 (3.2) |
| Completeness (%) | 98.61 (99.53) | 89.48 (90.89) | 96.78 (97.63) |
| Mean *I*/$\sigma$(*I*) | 7.83 (1.00) | 6.71 (1.49) | 6.96 (1.27) |
| Wilson *B*-factor | 40.44 | 63.59 | 54.2 |
| *R*<sub>merge</sub> | 0.166 (1.409) | 0.321 (1.216) | 0.165 (1.024) |
| *R*<sub>meas</sub> | 0.191 (1.633) | 0.398 (1.380) | 0.195 (1.219) |
| *R*<sub>pim</sub> | 0.099 (0.813) | 0.232 (0.634) | 0.101 (0.643) |
| CC1/2 | 0.98 (0.38) | 0.80 (0.53) | 0.98 (0.35) |
| Anomalous completeness (%) | 89.9 (90.6) | 78.1 (77.9) | 85.8 (84.8) |
| Anomalous multiplicity (%) | 1.8 (2.0) | 1.9 (2.3) | 1.6 (1.8) |

**Supplementary Table 5.** Distances to metal (Å) for coordinating atoms at the orthosteric site

|  | Outward open | Occluded |  | Inward open |  |  |
| --- | --- | --- | --- | --- | --- | --- |
| Coordinating atom | G223W•Mn<sup>2+</sup> | WT•Mn<sup>2+</sup> | A47W•Mn<sup>2+</sup> | M230A•Mn<sup>2+</sup> | D296A•Mn<sup>2+</sup><sup>e</sup> | WT•Cd<sup>2+</sup> |
| D56 Oδ1 | 2.4 | 2.7 | 2.6 | 2.7 | 2.7 |  |
| N59 Oδ1 | 3.2 | 2.4 | 2.5 | 2.6 | 2.4 | 2.9 |
| M230 Sδ | 3.0 | 2.8 | 2.8 |  | 2.7 | 3.4 |
| A227 O (Ca) |  | 2.5 | 2.2 | 2.4 | 2.2 | 2.8 |
| A53 O (Ca) | 2.4 | 2.2 | 2.1 |  |  |  |
| Y54 O (Ca) |  |  |  | 3.1 | 3.5 | 3.2 |
| water O (Q378)<sup>a</sup> | 2.7 | 2.3 | 2.2 | 2.5 | 2.3 | 2.9 |
| water O (M230A)<sup>b</sup> |  |  |  | 2.3 |  |  |
| water (Y54)<sup>c</sup> |  |  |  | 2.2 |  | 3.3 |
| water (N59)<sup>d</sup> | 2.6 |  |  |  |  |  |

<sup>a</sup> Conserved water coordinating central binding ion (water/metal) and Oε1 of Q378 (except in G223W•Mn<sup>2+</sup> where Q378 is far away to be in coordinating distance).
<sup>b</sup> Water replacing M230 in M230A•Mn<sup>2+</sup> structure.
<sup>c</sup> Water coordinating central metal ion and Ca carbonyl of Y54.
<sup>d</sup> Water coordinating central metal ion and Oδ1 and Nδ2 of N59 in G223W•Mn<sup>2+</sup>.
<sup>e</sup> Missing coordinating bonds due to low resolution of the D296A•Mn<sup>2+</sup> structure.

**Supplementary Table 6.**
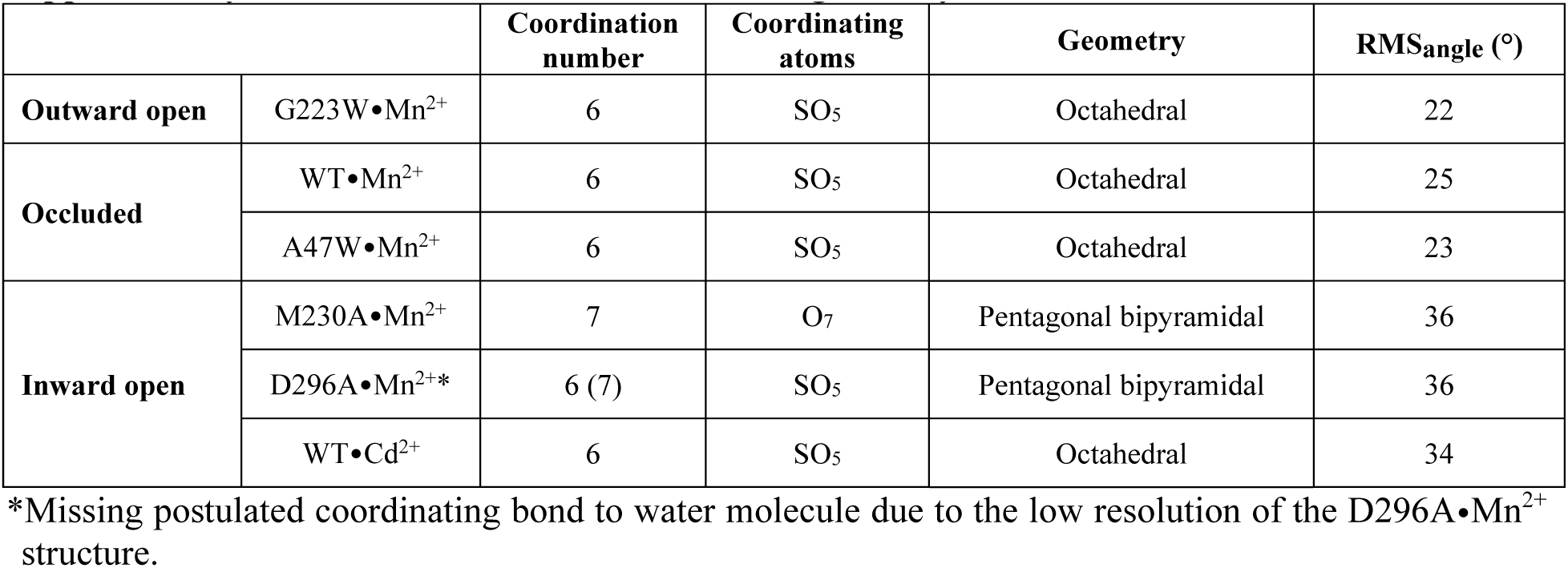
Coordination number and geometry of metal ions in the orthosteric site

**Supplementary Table 7.**
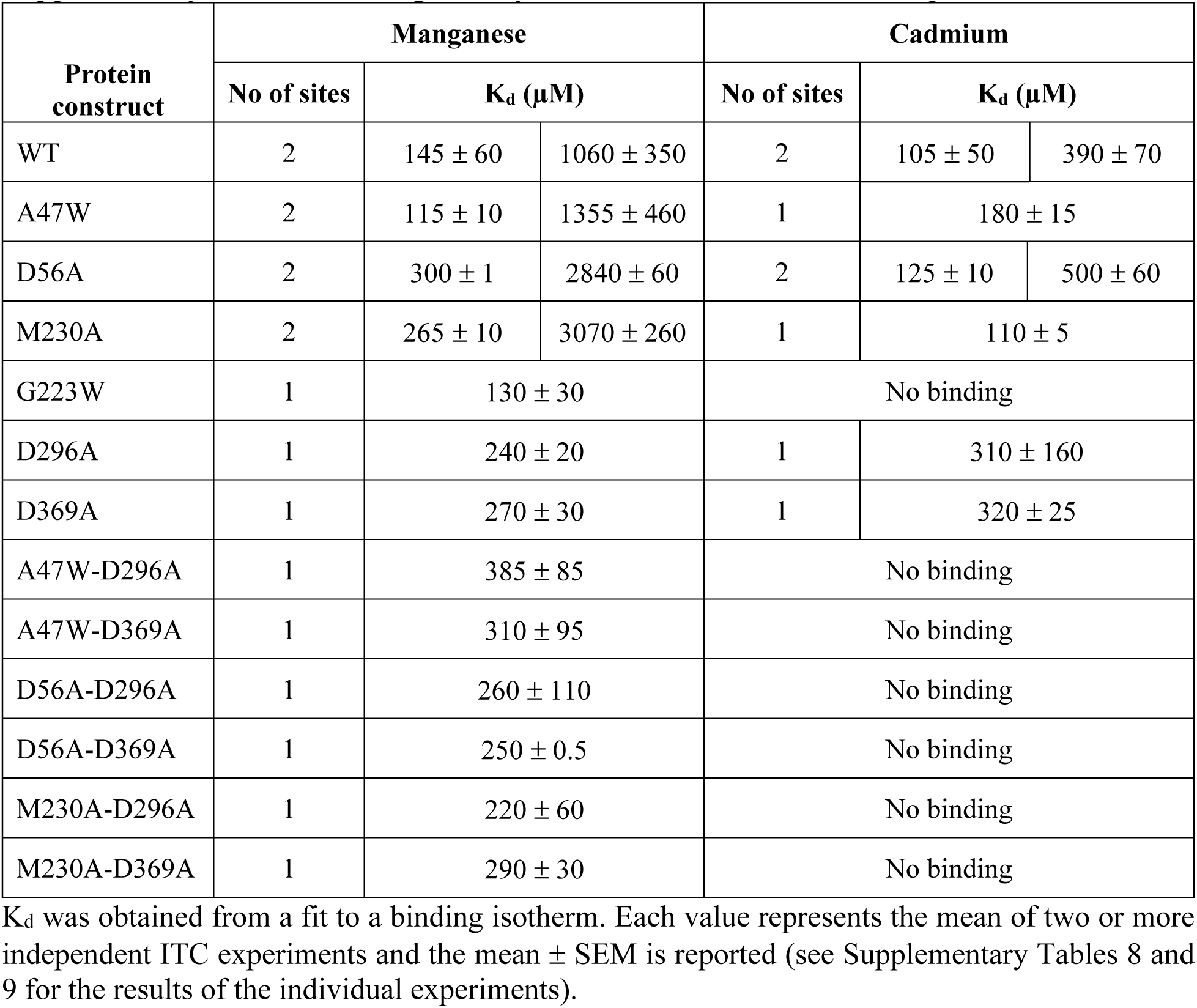
Binding affinity of metals to various DraNramp constructs

**Supplementary Table 8.** Thermodynamic parameters of Mn^2+^ binding to DraNramp constructs

| Protein construct | K<sub>d</sub> (μM) |  | ΔH (kcal mol<sup>-1</sup>) |  | -TΔS (kcal mol<sup>-1</sup>) |  | ΔG (kcal mol<sup>-1</sup>) |  |
| --- | --- | --- | --- | --- | --- | --- | --- | --- |
| WT | 191 | 840 | 2.7 | 5.7 | -7.7 | -9.8 | -5.0 | -4.1 |
|  | 224 | 1745 | 8.0 | 13.7 | -12.8 | -17.3 | -4.8 | -3.6 |
|  | 22 | 595 | 0.9 | 2.3 | -7.1 | -6.5 | -6.2 | -4.2 |
|  | **145 ± 60** | **1060 ± 350** | **3.9 ± 2.1** | **7.2 ± 3.4** | **-9.2 ± 1.8** | **-11.2 ± 3.2** | **-5.3 ± 0.3** | **-4.0 ± 0.2** |
| A47W | 123 | 1818 | 6.2 | 11.4 | -11.3 | -14.9 | -5.1 | -3.5 |
|  | 103 | 893 | 5.1 | 6.8 | -10.4 | -10.7 | -5.3 | -3.9 |
|  | **115 ± 10** | **1355 ± 460** | **5.6 ± 0.6** | **9.1 ± 2.3** | **-10.8 ± 0.4** | **-12.8 ± 2.1** | **-5.2 ± 0.2** | **-3.7 ± 0.2** |
| D56A | 298 | 2896 | 22.6 | 10.2 | -26.2 | -14.6 | -3.6 | -4.4 |
|  | 299 | 2780 | 9.7 | 15.6 | -14.0 | -18.5 | -4.3 | -2.9 |
|  | **298 ± 1** | **2840 ± 60** | **16.1 ± 6.5** | **12.9 ± 2.7** | **-20.1 ± 6.1** | **-16.5 ± 1.9** | **-4.0 ± 0.4** | **-3.6 ± 0.8** |
| M230A | 276 | 3333 | 3.0 | 14.9 | -7.7 | -18.2 | -4.7 | -3.3 |
|  | 254 | 2814 | 2.9 | 29.9 | -8.0 | -33.1 | -5.1 | -3.2 |
|  | **265 ± 10** | **3070 ± 260** | **3.0 ± 0.1** | **22.4 ± 7.5** | **-7.8 ± 0.1** | **-25.6 ± 7.4** | **-4.8 ± 0.1** | **-3.2 ± 0.1** |
| G223W | 104 |  | 10.5 |  | -14.0 |  | -3.5 |  |
|  | 163 |  | 5.8 |  | -10.1 |  | -4.3 |  |
|  | **130 ± 30** |  | **8.1 ± 2.3** |  | **-12.1 ± 1.9** |  | **-3.9 ± 0.4** |  |
| D296A | 202 |  | 8.9 |  | -13.4 |  | -4.5 |  |
|  | 278 |  | 14.5 |  | -18.7 |  | -4.2 |  |
|  | 229 |  | 14.3 |  | -18.2 |  | -3.9 |  |
|  | **240 ± 20** |  | **12.6 ± 1.8** |  | **-16.8 ± 1.7** |  | **-4.2 ± 0.1** |  |
| D369A | 252 |  | 14.5 |  | -18.8 |  | -4.3 |  |
|  | 235 |  | 14.7 |  | -18.8 |  | -4.1 |  |
|  | 225 |  | 8.9 |  | -13.4 |  | -4.5 |  |
|  | **270 ± 30** |  | **12.7 ± 1.9** |  | **-17.0 ± 1.8** |  | **-4.3 ± 0.1** |  |
| A47W-D296A | 299 |  | 6.2 |  | -10.7 |  | -4.5 |  |
|  | 471 |  | 13.1 |  | -17.6 |  | -4.5 |  |
|  | **385 ± 85** |  | **9.6 ± 3.5** |  | **-14.1 ± 3.4** |  | **-4.5 ± 0.0** |  |
| A47W-D369A | 403 |  | 6.5 |  | -11.0 |  | -4.5 |  |
|  | 213 |  | 7.6 |  | -12.5 |  | -4.9 |  |
|  | **310 ± 95** |  | **7.0 ± 0.1** |  | **-11.7 ± 0.7** |  | **-4.7 ± 0.6** |  |
| D56A-D296A | 427 |  | 10.0 |  | -14.6 |  | -4.6 |  |
|  | 286 |  | 6.2 |  | -11.0 |  | -4.8 |  |
|  | 55 |  | 2.1 |  | -7.7 |  | -5.6 |  |
|  | **260 ± 110** |  | **6.1 ± 2.3** |  | **-11.1 ± 1.9** |  | **-5.0 ± 0.4** |  |
| D56A-D369A | 249 |  | 3.9 |  | -8.9 |  | -5.0 |  |
|  | 250 |  | 3.2 |  | -7.4 |  | -4.2 |  |
|  | **250 ± 0.5** |  | **3.5 ± 0.4** |  | **-8.1 ± 0.7** |  | **-4.6 ± 0.3** |  |
| M230A-D296A | 321 |  | 7.1 |  | -11.6 |  | -4.5 |  |
|  | 214 |  | 0.5 |  | -5.4 |  | -4.9 |  |
|  | 115 |  | 3.3 |  | -8.6 |  | -5.3 |  |
|  | **220 ± 60** |  | **3.6 ± 1.9** |  | **-8.5 ± 1.8** |  | **-4.9 ± 0.1** |  |
| M230A-D369A | 319 |  | 7.4 |  | -11.6 |  | -4.2 |  |
|  | 257 |  | 11.4 |  | -15.5 |  | -4.1 |  |
|  | **290 ± 30** |  | **9.4 ± 2.0** |  | **-13.5 ± 1.9** |  | **-4.1 ± 0.1** |  |

K<sub>d</sub>, ΔH and ΔS were obtained from a fit to a binding isotherm, while ΔG was calculated from ΔG = ΔH - TΔS where T = 298 K. For each sample, the values for all runs are presented along with their mean ± SEM in bold. Grey cells correspond to values assigned to the external metal-binding site.

**Supplementary Table 9.** Thermodynamic parameters of Cd^2+^ binding to DraNramp constructs

| Protein construct | K<sub>d</sub> (μM) |  | ΔH (kcal mol<sup>-1</sup>) |  | -TΔS (kcal mol<sup>-1</sup>) |  | ΔG (kcal mol<sup>-1</sup>) |  |
| --- | --- | --- | --- | --- | --- | --- | --- | --- |
| WT | 85 | 245 | -4.3 | -2.6 | -1.2 | -2.1 | -5.5 | -4.7 |
|  | 35 | 429 | -1.6 | -2.5 | -4.2 | -2.1 | -5.8 | -4.6 |
|  | 195 | 485 | -3.5 | -6.5 | -1.5 | -1.8 | -5.0 | -8.3 |
|  | **105 ± 50** | **390 ± 70** | **-3.2 ± 0.8** | **-3.9 ± 1.3** | **-2.3 ± 0.95** | **-2.0 ± 0.09** | **-5.5 ± 0.1** | **-5.9 ± 1.2** |
| D56A | 115 | 563 | -5.8 | -6.8 | -0.3 | -1.5 | -6.1 | -8.3 |
|  | 133 | 437 | -2.5 | -4.3 | -1.2 | -1.8 | -3.7 | -6.1 |
|  | **125 ± 10** | **500 ± 60** | **-4.2 ± 1.6** | **-5.5 ± 1.3** | **-0.7 ± 0.4** | **-1.6 ± 0.1** | **-4.9 ± 1.2** | **-7.2 ± 1.2** |
| A47W | 163 |  | -4.8 |  | -2.9 |  | -7.7 |  |
|  | 191 |  | -4.9 |  | -2.9 |  | -7.8 |  |
|  | **180 ± 15** |  | **-4.9 ± 0.1** |  | **-2.9 ± 0.0** |  | **-7.7 ± 0.1** |  |
| M230A | 116 |  | -6.1 |  | -0.6 |  | -6.7 |  |
|  | 106 |  | -5.4 |  | -0.1 |  | -5.5 |  |
|  | **110 ± 5** |  | **-5.7 ± 0.3** |  | **-0.3 ± 0.3** |  | **-6.0 ± 0.6** |  |
| G223W | **No binding** |  |  |  |  |  |  |  |
| D296A | 122 |  | -3.5 |  | -1.8 |  | -5.3 |  |
|  | 171 |  | -2.3 |  | -2.7 |  | -5.0 |  |
|  | 637 |  | -3.3 |  | -0.9 |  | -4.2 |  |
|  | **310 ± 160** |  | **-3.0 ± 0.3** |  | **-1.8 ± 0.5** |  | **-4.8 ± 0.2** |  |
| D369A | 271 |  | -1.4 |  | -4.8 |  | -6.2 |  |
|  | 358 |  | -2.1 |  | -3.8 |  | -5.9 |  |
|  | 320 |  | -3.3 |  | -1.5 |  | -4.8 |  |
|  | **320 ± 25** |  | **-2.2 ± 0.5** |  | **-3.7 ± 0.8** |  | **-5.9 ± 0.3** |  |
| A47W-D296A | **No binding** |  |  |  |  |  |  |  |
| A47W-D369A | **No binding** |  |  |  |  |  |  |  |
| D56A-D296A | **No binding** |  |  |  |  |  |  |  |
| D56A-D369A | **No binding** |  |  |  |  |  |  |  |
| M230A-D296A | **No binding** |  |  |  |  |  |  |  |
| M230A-D369A | **No binding** |  |  |  |  |  |  |  |

K<sub>d</sub>, ΔH and ΔS were obtained from a fit to a binding isotherm, while ΔG was calculated from ΔG = ΔH - TΔS where T = 298 K. For each sample, the values for all runs are presented along with their mean ± SEM in bold. Grey cells correspond to values assigned to the external metal-binding site.

**Supplementary Table 10.** Summary of molecular dynamics simulations

| Simulation | Time (ns) | Starting structure | Number of atoms |
| --- | --- | --- | --- |
| S1a | 1031.95 | Outward-open, G223W•Mn<sup>2+</sup> (PDB ID 6BU5) with Mn<sup>2+</sup> removed and residue 223 mutated back to native glycine *in silico* | 104,624 |
| S1b | 617.55 |  |  |
| S2a | 769.80 | New WT•Mn<sup>2+</sup> inward-occluded structure, with Mn<sup>2+</sup> removed | 103,074 |
| S2b | 1022.85 |  |  |
| S3a | 1176.00 | New WT•Cd<sup>2+</sup> inward-open structure, with Cd<sup>2+</sup> removed | 103,372 |
| S3b | 808.80 |  |  |

**Supplementary Figure 1.**
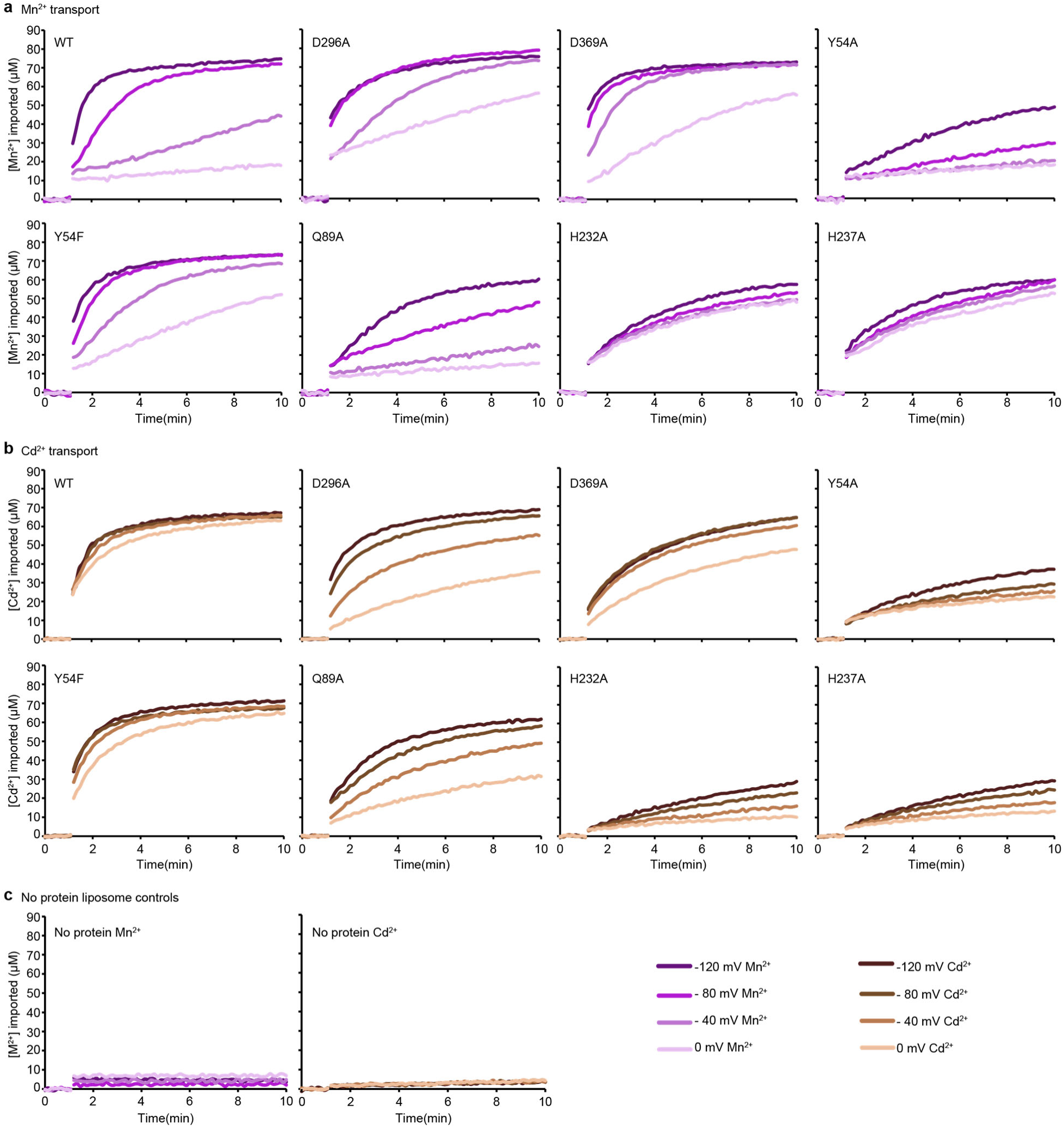
Representative time traces of metal ion uptake into proteoliposomes (n = 2-3) measured at four ΔΨ values for each DraNramp construct. (a,b) Mn^2+^ (a) and Cd^2+^ (b) uptake for WT and mutant DraNramp constructs. (c) Representative time traces of Mn^2+^(left) and Cd^2+^(right) uptake (n = 3) shows that no metal was imported into control liposomes. The initial metal uptake rates calculated from these time traces are in Figure 3c, Extended Data Figure 1c and Extended Data Figure 6f.

**Supplementary Figure 2.**
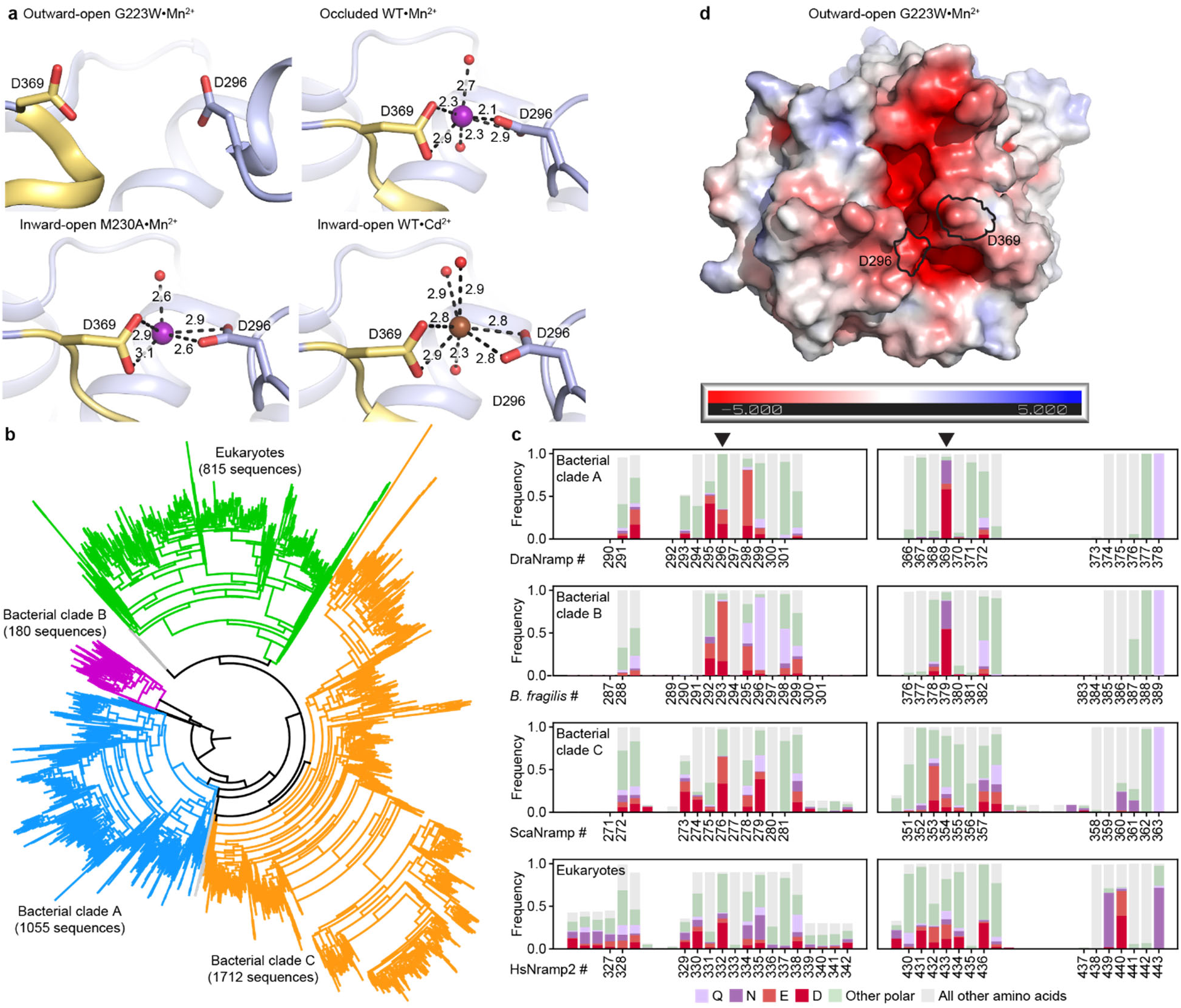
The external metal-binding site in DraNramp is somewhat conserved in clade A homologs, but poorly conserved across all Nramps. (a) External site architecture in the G223W•Mn^2+^ (outward-open), WT•Mn^2+^ (occluded), M230A•Mn^2+^ and WT•Cd^2+^ (inward-open) structures showing positions of D296 and D369 across conformations, illustrating that the two residues are farther apart in the outward-open structure and cannot bind metal. Metal-coordinating distances are listed in Å. (b) A maximum likelihood phylogenetic tree illustrating evolutionary divergence of the Nramp family into several major clades for prokaryotes (clades A, B and C) and eukaryotes. (c) Frequencies of acidic and other polar amino acids in the loop regions surrounding D296 (left; loop preceding EH2) and D369 (right; loop preceding TM10) across phylogenetic clades, based on the sequence alignment used to build the tree in panel b (bacterial clade A, DraNramp numbering; bacterial clade B; *Bacteroides fragilis* MntH numbering; bacterial clade C; ScaNramp numbering; eukaryotic clade, human Nramp2 numbering). Across all clades, these two external loops have a high concentration of acidic amino acids. However, the exact positions of acidic residues observed in DraNramp—296 and 369 (arrowheads)—are not highly conserved except 369 in bacterial clades A and B, which is an aspartate, asparagine, or glutamate in most sequences (92.1% and 87.8% in clades A and B, respectively) (d) APBS^2^-generated electrostatic surface potential of the outward-open structure viewed from the extracellular side illustrates that D296 and D369 contribute to a funnel of negative charge leading into the orthosteric binding site.

**Supplementary Figure 3.**
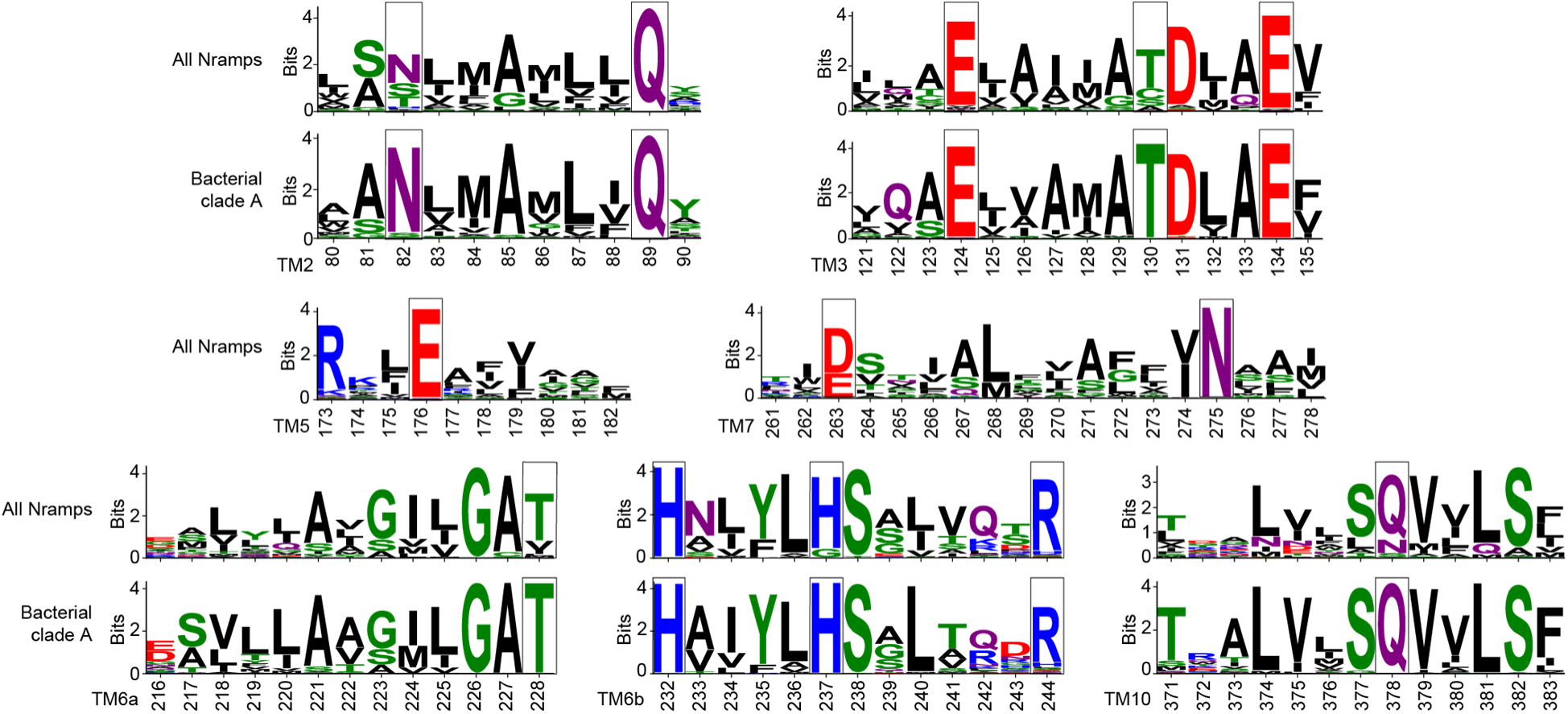
Sequence logos showing conservation of polar residues important for opening and closing the gates during Mn^2+^ transport (highlighted in black boxes). The alignment contains 3762 Nramp sequences, including 1055 bacterial clade A Nramps. Most residues that are less conserved in all Nramps are completely conserved in clade A to which DraNramp belongs. Residue coloring is based on the WebLogo scheme for amino acid chemistry^3^.

**Supplementary Figure 4.**
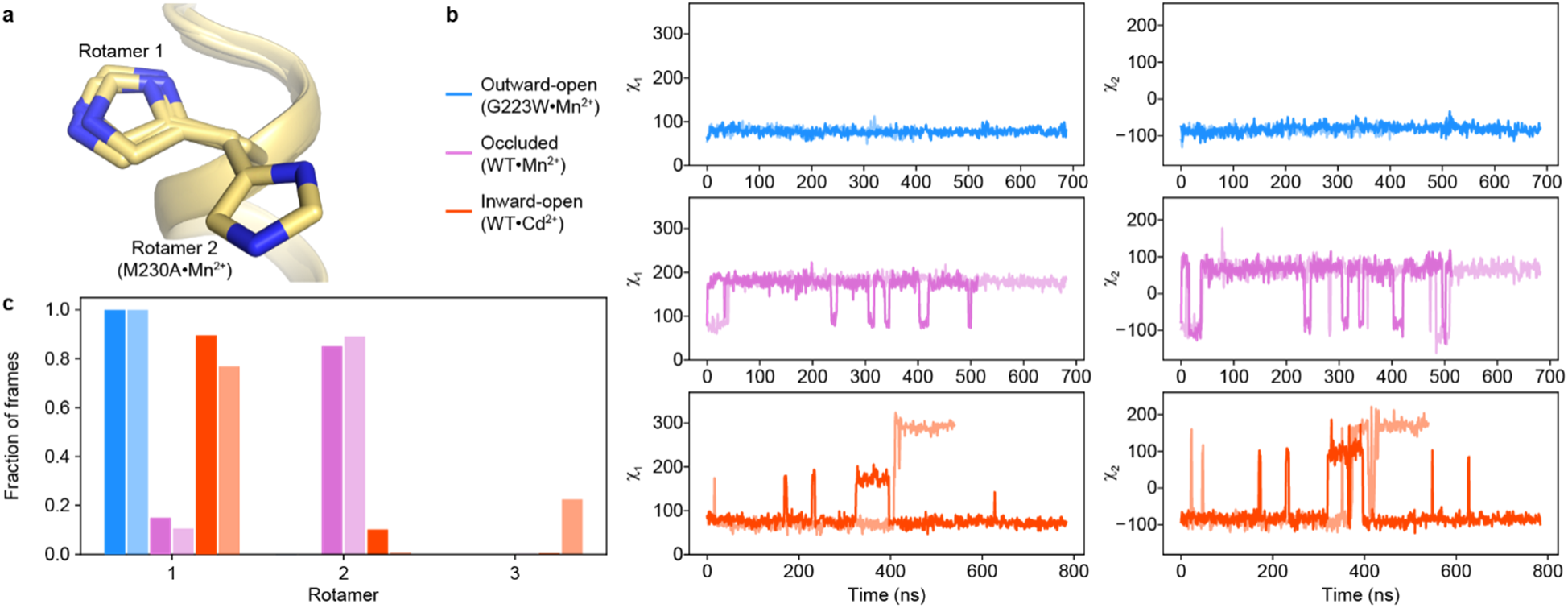
The H232 rotamer is fixed in the outward-open state but dynamic in the occluded and inward-open states. (a) H232 takes on rotamer 1 in all structures, except in the M230A structure where it takes on rotamer 2. (b) Time-course analyses of the χ_1_ and χ_2_ angles of the H232 sidechain during the MD simulations starting in each of the three major conformations, showing replicates in different shades. Interestingly, the simulations in the occluded state show H232 rapidly switching to rotamer 2 and primarily occupying the rotamer 2 state throughout the simulation. In contrast, H232 is fixed in rotamer 1 throughout the outward-open simulation and shows the most flexibility in the inward-open state. (c) Distribution of H232 rotamer states across each MD simulation.

